# Marine food webs are more complex but less stable in sub-Antarctic than in Antarctic regions

**DOI:** 10.1101/2021.04.12.439560

**Authors:** Iara Diamela Rodriguez, Tomás Ignacio Marina, Irene Ruth Schloss, Leonardo Ariel Saravia

## Abstract

Food web structure plays an important role in determining ecosystem stability against perturbations. High-latitude marine ecosystems are being affected by environmental stressors and biological invasions. In the West Antarctic Peninsula these transformations are mainly driven by climate change, while in the sub-Antarctic region by anthropogenic activities. Understanding the differences between these areas is necessary to monitor the changes that are expected to occur in the upcoming decades. Here, we compared the structure and stability of Antarctic (Potter Cove) and sub-Antarctic (Beagle Channel) marine food webs. We compiled species trophic interactions (predator-prey) and calculated complexity, structure and stability metrics. Even if both food webs presented the same connectance, we found important differences between them. The Beagle Channel food web is more complex, but less stable and sensitive to the loss of its most connected species, while the Potter Cove food web presented lower complexity and greater stability against perturbations.

## 1 Introduction

Food webs represent the trophic interactions (predator-prey) that occur in an ecosystem, aiding to understand the flows of energy and matter among organisms. Trophic interactions are key drivers of ecosystems structure, function and stability (Dunne 2006). The occurrence, intensity and frequency of these interactions can and are being altered by climate change and anthropogenic stressors, by modifying the species patterns of distribution and abundance (Blois et al. 2013).

Despite that food web structure shows some generalities across habitats (Mora et al., 2018), it is expected that particular environmental and biogeographic conditions will shape network architecture in different ways (Song and Saavedra 2020). For instance, temperature was proposed to be positively associated with food web complexity (connectance), number of trophic levels and omnivory (Gibert 2019; Kortsch et al. 2018); habitat heterogeneity promotes food web complexity in terms of number of species and trophic interactions, trophic levels and omnivory (Kortsch et al. 2018). Furthermore, ecosystems with higher productivity can sustain more species (Duffy, Godwin, and Cardinale 2017) and longer food chains (Young et al. 2013).

High-latitude marine ecosystems are being affected by drastic environmental and ecological transformations, driven by climate change and anthropogenic activities (Clarke and Harris 2003; Hoegh-Guldberg and Bruno 2010; Meredith et al. 2019). The West Antarctic Peninsula (WAP) had the highest heating rates recorded worldwide in the past half-century (Steig et al. 2009; Turner et al. 2014). It is considered that this area is undergoing a transition from a cold-dry polar-type to a warm-humid sub-Antarctic-type climate (Montes-Hugo et al. 2009), while human activities intensify (McCarthy et al., 2019; IAATO, 2019). As a consequence, physical and chemical changes (e.g., glacier retreat, increased sediment input, seasonal sea-ice reduction, surface salinity decrease) and biological responses are increasingly being reported (Cook et al. 2005; Ducklow et al. 2013; Fuentes et al. 2016; Schloss et al. 2012). Changes in communities’ composition, species distribution and abundance (Lagger et al. 2017; Pasotti et al. 2015; Sahade et al. 2015) support such climate transition hypothesis.

At the southernmost tip of South America, it is located the Beagle Channel, a sub-Antarctic drowned glacial valley. It is the closest coastal area to the WAP and shares a relatively recent biogeographical history, from around 30 Ma, when the Antarctic Circumpolar Current was established (Barker and Thomas 2004). As a result, these areas became two distinct biogeographical regions, with different environmental characteristics and biodiversity (Griffiths and Waller 2016). Nowadays, the disturbances that affect such ecosystems have different origins. The Beagle Channel is mainly threatened by anthropogenic pressures: increasing levels of pollution (urban sewage, industrial activities, shipping traffic and tourism) (Amin, Ferrer, and Marcovecchio 1996; Gil et al. 2011), the introduction of exotic salmon species for aquaculture (Fernández et al. 2010; Nardi et al. 2019) and the fishing pressure of economically important species (Lovrich, 1997). Although there is no clear evidence that this area is currently being affected by climate change, marine temperatures are projected to globally increase in the next 100 years (IPCC, 2013). Species distributions are expected to move poleward with warming, to maintain their preferred temperature range (Hickling et al. 2006). There are already reports of sub-Antarctic alien species in Potter Cove and the WAP (Cárdenas et al. 2020; Fraser et al. 2018) and many have been proposed to have the potential to invade Antarctica (Diez and Lovrich 2010; Hughes et al. 2020; López-Farrán et al. 2021). Understanding the differences (and similarities) between sub-Antarctic and Antarctic areas is necessary to monitor the changes that are occurring and are expected to increase in the upcoming decades (Griffiths, Meijers, and Bracegirdle 2017; Gutt et al. 2015). Developing models help us predict these ecosystems’ responses.

In this study, we use a network approach to explore and compare the structure and stability of Antarctic (Potter Cove, South Shetland Islands, WAP) and sub-Antarctic (Beagle Channel, Tierra del Fuego) food webs. We hypothesize that the relatively warmer, more productive Beagle Channel (Amin et al. 2011) will present a more complex food web. Consequently, the food web is expected to present a lower stability to cope with perturbations (May 1973). On the other hand, the strongly seasonal, ice-covered and less productive (Schloss et al. 2012) Potter Cove will present a less complex and more stable food web.

## 2 Materials and methods

### 2.1 Study sites

The Beagle Channel (Figure 1a) connects the Pacific and Atlantic Oceans and is about 240 km long and 5 km wide, at its narrowest point. To facilitate the comparison, we selected a study area that comprises two distinguishable coastal sites: Lapataia Bay (54° 51’S, 68° 34’W), located within the Tierra del Fuego National Park, with a maximum depth of 20 m and rocky bottoms with abundant macroalgae; and Ushuaia Bay (54° 49’S, 68° 19’W), where the city of Ushuaia is located, 140 m deep and with a soft substrate with small rocks and shells (Balestrini, Manzella, and Lovrich 1998). The coastline of the Beagle Channel is characterized by the presence of giant kelp (*Macrocystis pyrifera)* forests. Due to their complex morphology, kelps act as ecosystem engineers, providing refuge from predation and a habitat for prey, and altering water conditions (Adami and Gordillo 1999; Bruno et al. 2018; Graham, Vásquez, and Buschmann 2007).

**Figure 1.**
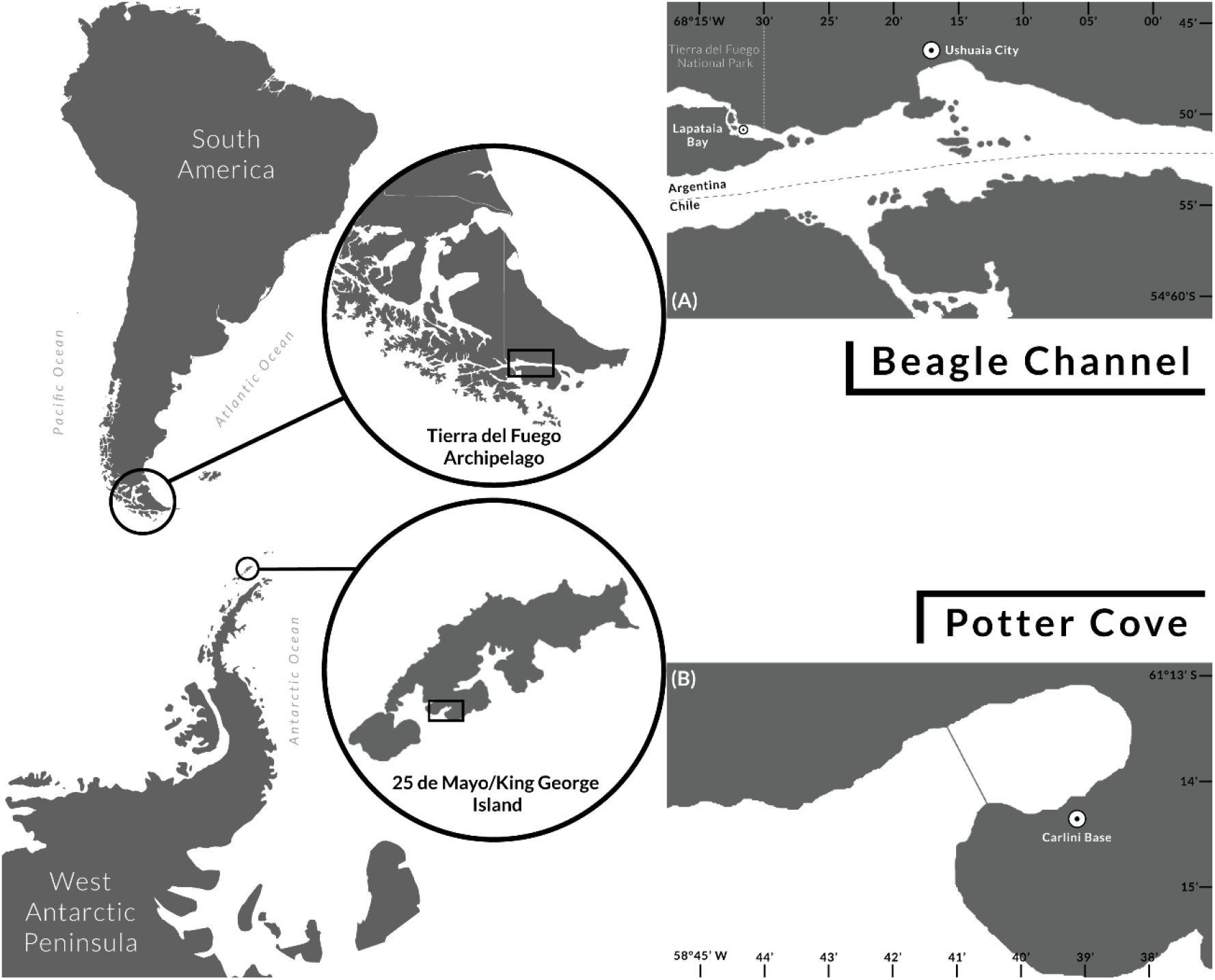
Map of **(a)** Beagle Channel at Tierra del Fuego Archipelago, South America and **(b)** Potter Cove at 25 de Mayo/King George Island, West Antarctic Peninsula. (Map originally taken from FreeVectorMaps.com https://freevectormaps.com)

Potter Cove is a fjord-like environment located at 25 de Mayo/King George Island (62° 14’S, 58° 38’W, South Shetland Islands) on the West Antarctic Peninsula (Figure 1b). It has an area of 7 km^2^ (4 km long and 2.5 km wide). A shallow sill (<30 m) separates the cove into an inner, shallower (<50 m) part characterized by soft sediments inhabited by benthic filter feeder species from a deeper (∼100 m) outer section, with rocky bottom, colonized by a large biomass of macroalgae. Due to their high-latitude location, marine Antarctic ecosystems are extremely variable due to the strong seasonality in the photoperiod length. Winter reduction in irradiance and temperature controls the dynamics of environmental variables, such as incident radiation, sea-ice extent, mixing layer depth, water column particulate matter and nutrients concentration.

Water temperature, in Potter Cove, ranges from −2 to −1°C during winter and 0.3 to 2.1°C during summer (Schloss et al. 2012). In the Beagle Channel temperature ranges from 4.2 - 4.3°C in winter, to 8.9 - 9.8°C in summer (Balestrini, Manzella, and Lovrich 1998). Both sites have high availability of nutrients, meaning that the main limiting factor conditioning primary production is solar irradiance (Amin et al. 2011; Schloss, Ferreyra, and Ruiz-Pino 2002). During summer, sediment runoff into the water column increases due to the input of glacier meltwater at both sites.

### 2.2 Food web assembly and topology

We collected and compiled trophic interactions (prey-predator) of species present in each site. For Potter Cove, we updated the food web published by Marina et al. (2018), maintaining the same criteria they used in their assembly: consider only trophic links confirmed by gut content studies and/or field observations; experimental and biomarkers (isotopes and fatty acids) studies were not taken into account; mammals, seabirds and pelagic fish were not considered since their contribution is not relevant to the flow of matter and energy in the cove (Barrera Oro and Casaux 2008). Most data were collected during austral summer months, when most research campaigns are carried out. More detailed information on Potter Cove food web assembly can be found in Marina et al. (2018).

Little data on species diets was available for the chosen sites of the Beagle Channel, as compared to Potter Cove. We decided to include information from biomarkers studies (Riccialdelli et al., 2017) corroborating the diet information with the bibliography found on them. When there was insufficient species diet information, we included trophic links reported from nearby sub-Antarctic ecosystems, such as the Magellan region and Navarino Island on the Beagle Channel (Castilla 1985; Díaz 2016; Moreno and Jara 1984). We also took into account diet information collected from seasons other than summer, since in Beagle Channel trophic interactions do not fluctuate significantly during the year, but rather species abundance does (Almandoz et al. 2011; Adami and Gordillo 1999; Aguirre et al. 2012). Mammals and seabirds were also not considered, although we included pelagic fish given their abundance and importance as prey in Beagle Channel (Riccialdelli et al., 2020).

The trophic network was defined by an adjacency matrix (A) of pairwise interactions, in which each element a_ij_ = 1 when the j-species preyed on the i-species, and a_ij_ = 0 otherwise. From this matrix one can obtain a directed graph with L trophic links connecting S nodes or trophic species. Trophic species can correspond to: biological species groups, taxonomic groups above the species level due to lack of diet resolution (genus, family), organisms that share the same sets of predators and prey, and non-living compartments of matter (e.g., detritus, necromass). Henceforth, we will use the term “species” as synonymous with “trophic species” (Briand and Cohen 1984).

We described the network structure and complexity with metrics that are widely used in food web studies (Delmas et al. 2018; Landi et al. 2018; Montoya, Pimm, and Solé 2006), such as link density, connectance, percentage of basal/intermediate/top species, mean trophic level and omnivory (Table 1).

**Table 1.**
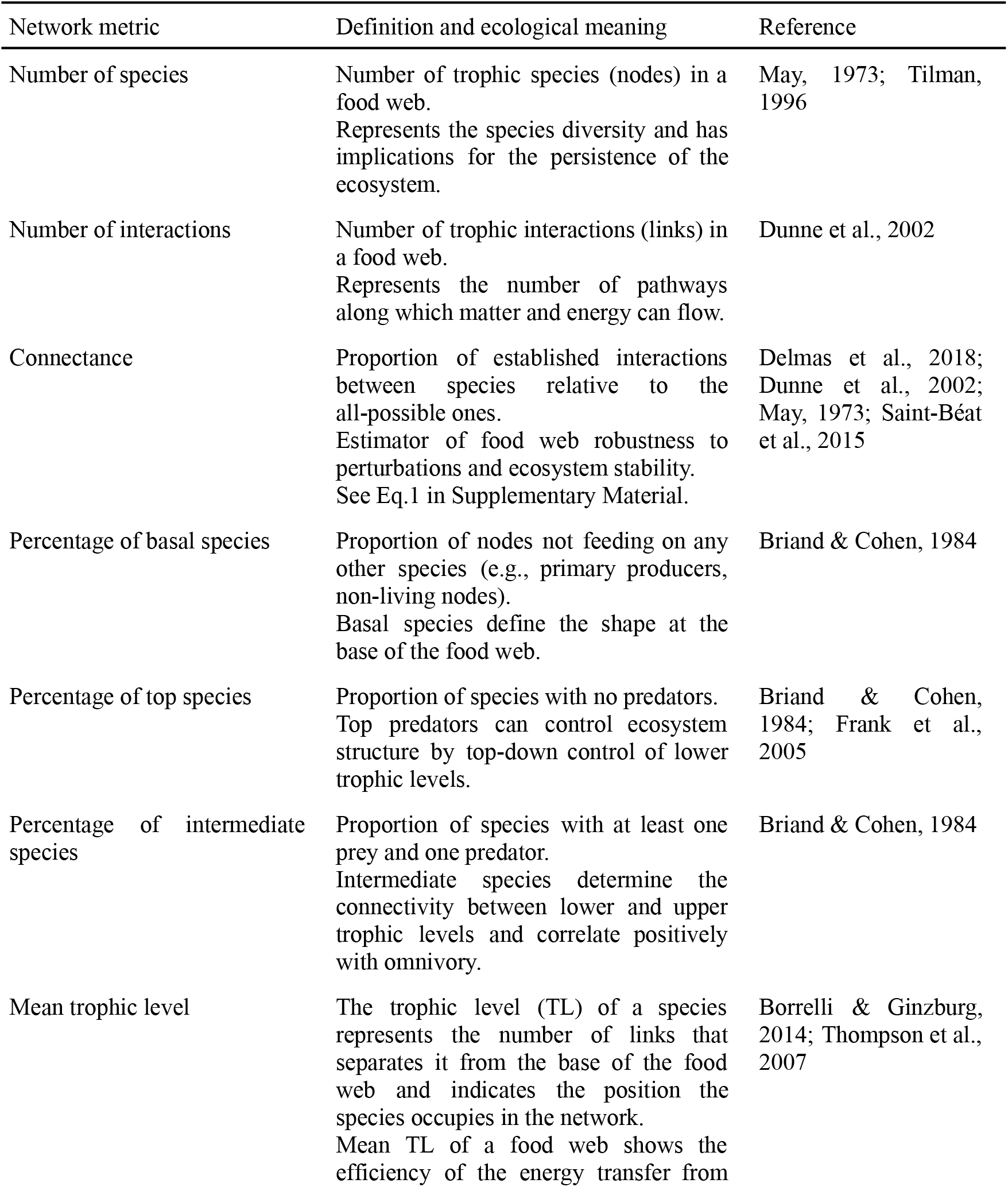
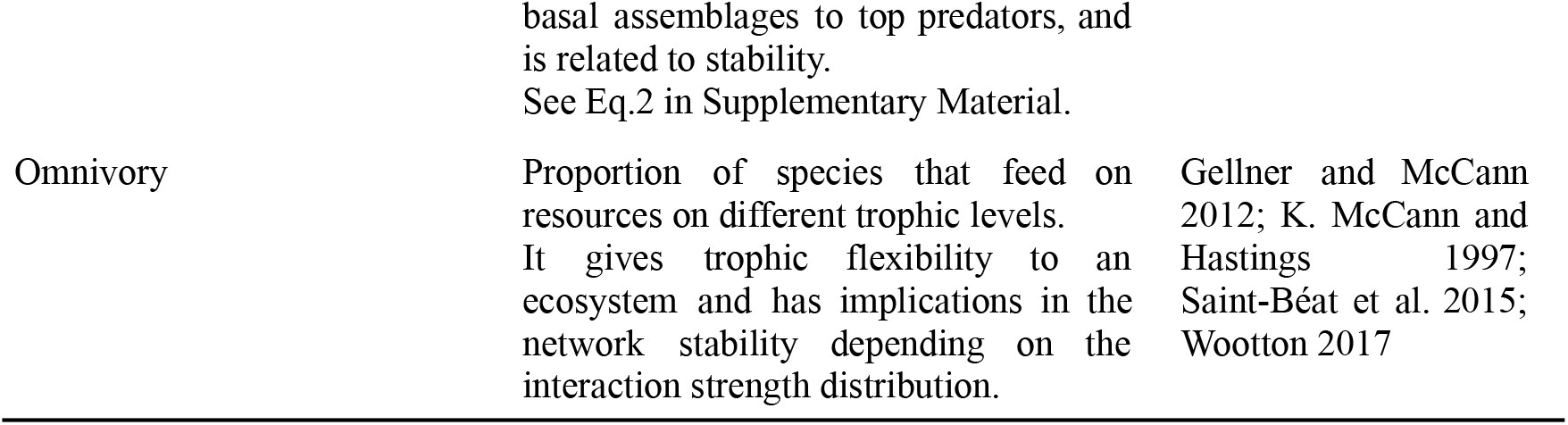
List of network metrics analyzed, definitions, and relevant ecological implication related to food web structure and complexity.

In addition, as a summary of the networks’ topology, we studied how trophic links were distributed among all species in the networks, the so-called cumulative degree distribution. For this purpose, node degree was calculated as the sum of all in-(number of prey) and out- (number of predators) trophic interactions. Then we fitted the cumulative degree distribution to the following models: exponential, log normal, poisson, power-law, power-law with exponential cutoff and uniform. Model fit was done using maximum likelihood (Mccallum 2008) and model selection was performed by the Akaike Information Criterion corrected for small sample size (Burnham and Anderson 2002).

### 2.3 Modularity, species roles and traits

Food webs tend to naturally organize in non-random, modular patterns (Grilli, Rogers, and Allesina 2016). This means that groups of prey and predators interact more strongly with each other, than with species belonging to other groups. Modularity measures how strongly these subgroups of species, called modules, interact with each other compared to species from other modules (See Eq.3 in Supplementary Material). Modular organization is positively associated with stability and enhances the persistence of a food web, since perturbations can be retained within modules, constraining the spreading to the rest of the network (Stouffer and Bascompte 2011).

Species can play different roles with respect to modularity, according to the pattern of trophic links they have within their own module and/or across modules. Nodes with the same role are expected to have similar topological properties (Guimera 2005; Kortsch et al. 2015). To evaluate species’ role similarity/dissimilarity between Potter Cove and Beagle Channel, we computed the topological role for each species, classified as: *module hub*, species with a relatively high number of links, but most within its own module; *module specialist*, species with relatively few links and most within its own module; *module connector*, species with relatively few links mainly between modules and *network connector*, species with high connectivity between and within modules (See Eq.4 in Supplementary Material). We combined topological roles with each species trophic level and module membership in one figure to provide an integrated visualization of these food web properties.

We also collected species’ biology and feeding behavior information and classified them regarding habitat (pelagic, benthopelagic and benthic) and functional group (non-living, basal taxa, zooplankton, benthos and fish), in order to determine if module affiliation was associated with these traits, as observed in other marine food webs (Kortsch et al. 2015; Rezende et al. 2009). To determine if the proportions of species per traits changed across modules and among trait levels between food webs, we used Pearson’s Chi-squared test and plotted the number of species habitat and functional group affiliation within each module and the percentage of species per trait level.

### 2.4 Quasi Sign-Stability (QSS)

Stability is a multidimensional concept (Donohue et al. 2013) and can be measured in different ways. Traditionally, it is conceived as the ability of an ecosystem to maintain its state over time, against external and internal forces that drive it away from that state (Saint-Béat et al. 2015). To evaluate and compare network stability between the Potter Cove and Beagle Channel food webs, we used a variation of the Quasi Sign-Stability metric (QSS) (Allesina and Pascual 2008). To calculate QSS, we built the community matrix (i.e., the jacobian) based on the adjacency matrix A and randomized the magnitude of the elements of the community matrix, following an asymmetrical distribution. This means that the impact of predators on prey is greater than the impact of prey on predators (Borrelli and Ginzburg 2014). QSS is typically considered as the proportion of community matrices with the maximal negative eigenvalue. In our variation, we calculated it as the mean of the maximum eigenvalue for the random community matrices for easier analysis and visualization. This metric is directly related with network local stability, which can reveal the amplification of small perturbations near the equilibrium point (Allesina and Pascual 2008); values closer to zero indicate a more stable food web.

### 2.5 Food web structure and stability comparison using a randomization algorithm

In order to perform a statistically robust comparison between food webs, we used the Strona Curveball algorithm (Strona et al. 2014) to generate an ecological meaningful distribution of the metrics (Cordone et al. 2020; Kéfi et al. 2016). It randomizes the network structure maintaining the number of prey and predators for each species, meaning that species have the same degree, but allow them to interact with different species than in the original network. We performed 1000 network randomizations for each Potter Cove and Beagle Channel food web and calculated structure and stability metrics (mean trophic level, omnivory, modularity, QSS). Complexity metrics (number of species, link density, connectance) were not calculated since they do not vary due to the algorithm restrictions. If empirical values for each metric were within the distribution of the randomized simulated food webs, we considered that simulations fitted the empirical values enabling network comparison. Then, we calculated the 95% confidence interval and compared the distributions obtained for each metric using the two-sided Kolmogorov-Smirnov test (Massey 1951).

### 2.6 Data analysis

All analyses, simulations and graphs were performed in R version 4.0.3 (R Development Core Team 2020), using the ‘PoweRlaw*’* R package to fit distributions (Gillespie 2015) and the ‘multiweb’ R package to calculate all network metrics and food web simulations (Saravia et al. 2019).

## 3 Results

### 3.1 Food web topology

Potter Cove and Beagle Channel food webs differed with regard to complexity and structural properties. Potter Cove food web included a total of 110 preys and predators, while 145 species characterized the Beagle Channel food web. Both networks had 3 non-living nodes (fresh detritus, aged detritus and necromass). Beagle Channel presented nearly twice the amount of total feeding interactions (1115) than Potter Cove (649), but the same connectance value (0.05). It should be noted that, despite having different number of species, both food webs presented a very similar species distribution in basal (27% Potter, 25% Beagle) and top positions (6% Potter, 8% Beagle), being most of them intermediate species (67% for both ecosystems) (Table 2). Beagle Channel food web showed a higher mean trophic level (2.3) and percentage of omnivory (55%) than Potter Cove (2.2 mean trophic level and 46% omnivory) (Figure 2a, b).

**Table 2.** Network metric values for Potter Cove and Beagle Channel food webs.

| Network metric | Potter Cove | Beagle Channel |
| --- | --- | --- |
| Number of species | 110 | 145 |
| Number of interactions | 649 | 1115 |
| Connectance | 0.05 | 0.05 |
| Basal species (%) | 27 | 25 |
| Top species (%) | 6 | 8 |
| Intermediate species (%) | 67 | 67 |

**Figure 2.**
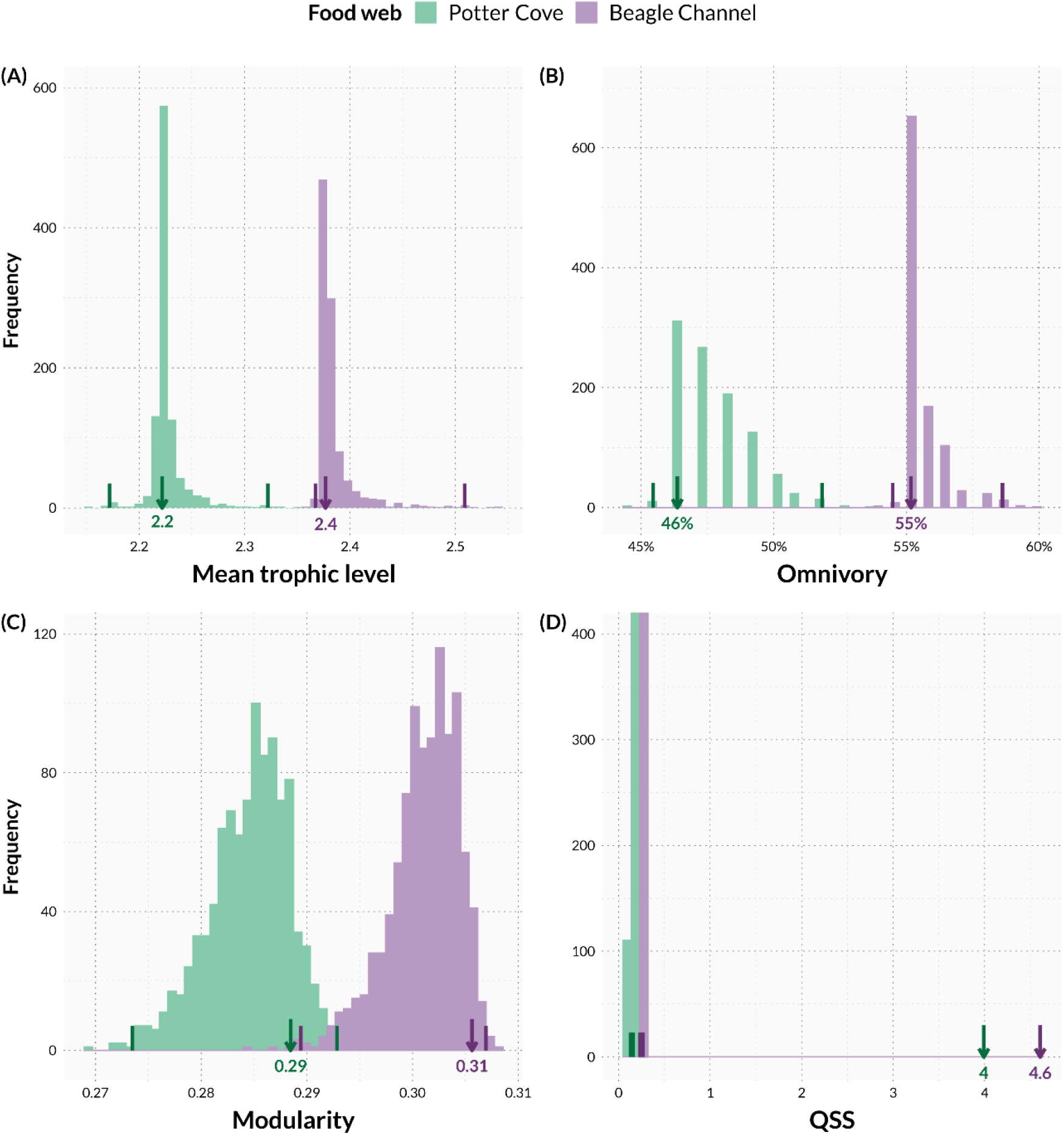
Randomization algorithm of food web structure and stability. **(a)** Mean trophic level and **(b)** omnivory, **(c)** modularity and **(d)** QSS histograms of simulated networks performed for Potter Cove (green) and Beagle Channel (purple) food webs. Empirical metric values are represented by arrows and the value is shown below the arrow. Small darker colored bars are randomizations’ lower and upper 95% confidence intervals. We found statistical differences (p<0.05) between food webs in all the analyzed metrics.

The cumulative degree distribution analysis showed strongly right-skewed degree distributions for both food webs, with most of the species with few links, and few species with many links. The best fit were the exponential model and the power-law with exponential cutoff for Potter Cove and Beagle Channel food webs, respectively, according to the AICc analysis (Supplementary Material Table S1). The power-law with exponential cutoff distribution is less steep than the exponential, which translates in Beagle Channel food web having more species with higher degree than Potter Cove’s.

### 3.2 Modularity, species roles and traits

Species topological role analysis showed that in both food webs most species are module connectors or specialists (species with few links, between modules or within its own module), with no module hub (species with many links within its own module) and only one network connector (species with many links between modules), but with different trophic position: the top predator (no species preying on it), black rockcod (*Notothenia coriiceps*) for Potter Cove, with a trophic level = 3.0; and the squat lobster (*Munida gregaria)* for Beagle Channel, with a trophic level = 2.4 (Figure 4).

**Figure 3.**
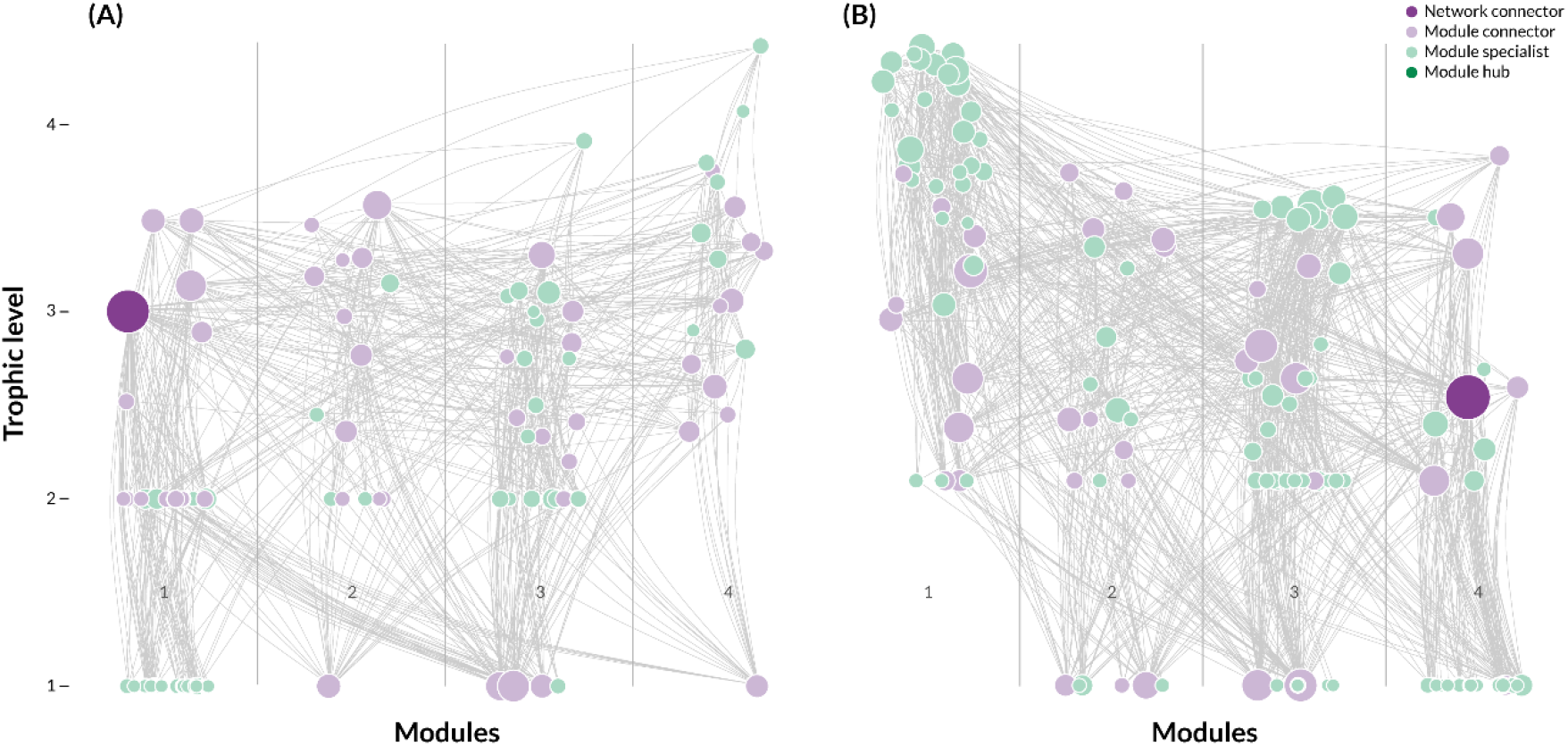
Food webs. Graphic representation of **(a)** Potter Cove and **(b)** Beagle Channel food webs. Each circle (node) represents a trophic species and the lines (links) represent feeding interactions between species. The vertical position of the nodes indicates the trophic position of a species and the horizontal position indicates the module affiliation of a species. The size of the nodes is proportional to the degree (number of links) of a species. The color of the node indicates species’ topological role: dark purple=network connector (species with high connectivity between and within modules), light purple=module connector (species with few links mostly between modules), light green=module specialist (species with few links within its own module), dark green=module hub (species with high number of links mostly within its own module).

**Figure 4.**
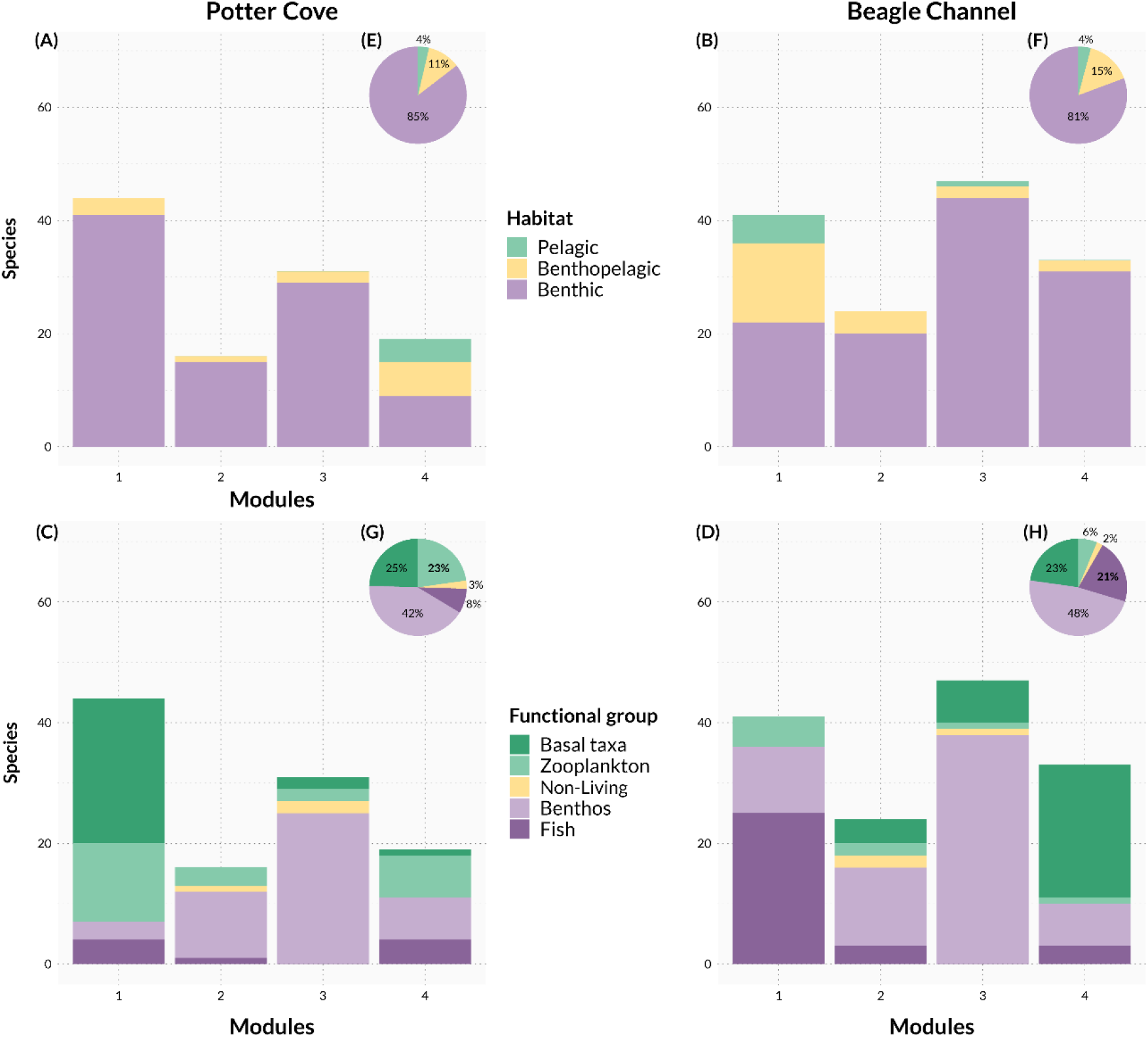
Food webs modules vs traits. Bar plot of the number of species within each module of **(a)** Potter Cove and **(b)** Beagle Channel colored by habitat: green=pelagic, yellow=benthopelagic, purple=benthic; significant differences were found between them (Potter Cove: Chi-squared = 34.43, p-value = <0.01. Beagle Channel: Chi-squared = 33.03, p-value = <0.01). Bar plots of **(c)** Potter Cove and **(d)** Beagle Channel food webs showing the frequency of species functional group affiliation: dark green=basal taxa, light green=zooplankton, yellow=non-living, light purple=benthos, dark purple=fish; significant differences were found between them (Potter Cove: Chi-squared = 76.53, p-value = <0.01. Beagle Channel: Chi-squared = 115.15, p-value = <0.01). Pie charts of species percentage within each type of habitat for **(e)** Potter Cove and **(f)** Beagle Channel; and functional groups for **(g)** Potter Cove and **(h)** Beagle Channel; significant differences were found only in the percentage of zooplankton and fish functional groups (Zooplankton: Chi-squared = 9.43, p-value = <0.01. Fish: Chi-squared = 5.89, p-value = 0.01).

Despite having significantly different modularity (which indicates how strongly the modules are tied together) (Figure 2f), the modularity analysis divided both food webs into four distinct modules (Figure 4). We found that the proportion of species’ traits (habitat and functional group) changed significantly across food webs’ modules. Another interesting fact is that the Beagle Channel food web had significantly three times more fish species (31) than Potter Cove (9), but presented a lower zooplankton ratio (0.23 Potter Cove, 0.06 Beagle Channel) (Figure 5g, h) due to lower taxonomic resolution for this last functional group.

A detailed list of all species, their module affiliation, role as network connector, module connector, module specialist and module hub, functional group, habitat use, degree and trophic level can be found in Supplementary Material Table S2 for Potter Cove and Table S3 for Beagle Channel.

### 3.3 Food web structure and stability comparison with a randomization algorithm

All the analyzed metrics of structure and stability showed statistical differences (p<0.01, Supplementary Material Table S4) between sites. An important remark is that all empirical values for the metrics fell within the distributions of the randomized simulated networks. Beagle Channel food web presented higher values for all of the analyzed metrics than Potter Cove (Figure 2).

## 4 Discussion

The comparison presented here shows that Potter Cove and Beagle Channel food webs have different structure, with important consequences for their stability. As we hypothesized, the sub-Antarctic food web is more complex, but less locally stable, and the Antarctic one exhibits lower complexity, which appears to provide stability in the face of perturbations.

### 4.1 Food web differences

The Beagle Channel food web displayed a higher number of trophic species (mainly due to higher fish richness), almost twice more links, higher mean trophic level and omnivory, meaning a more complex structure, as we hypothesized. More productive and larger ecosystems, like Beagle Channel, in comparison with Potter Cove, are expected to promote food web complexity. They can sustain a higher number of species and longer food chains, which correlates positively with omnivory, since species have a higher probability to encounter prey in different habitats and in a wider range of trophic positions (Kortsch et al. 2018; Thompson et al. 2007).

The different cumulative degree distributions show that the Beagle Channel presents a higher number of generalist species than the Potter Cove food web, supported by a greater proportion of omnivores, mainly fish. In contrast, the Antarctic coastal marine food web is known to contain many specialized benthic fish species (Barrera-Oro 2002). The distribution of species in the food web can also determine the vulnerability of the system to the loss of the most connected species. The power-law distribution presented by the Beagle Channel network suggests that this ecosystem is highly sensitive to perturbations affecting the species with highest degree, potentially driving the system to collapse (Albert, Jeong, and Barabási 2000; Estrada 2007). The emerging interest on the squat lobster (*Munida gregaria*), the most connected prey and abundant species (Arntz and Gorny 1994), might present a serious threat to the stability of this sub-Antarctic food web without a proper management (Tapella et al. 2002).

In theory, exponential networks such as Potter Cove’s, can be catastrophically fragmented by random removal of nodes (Albert, Jeong, and Barabási 2000). However, Cordone et al., (2018, 2020) simulated species extinction for Potter Cove and found no food web collapse, suggesting an apparent robustness to species loss. Our stability analysis supports their conclusion, as Potter Cove food web displayed significantly higher stability (lower QSS), meaning that it has a higher probability to recover after a perturbation, such as a local loss of a species, than the Beagle Channel food web.

The lower stability of the Beagle Channel food web, in comparison with Potter Cove, might be attributed to its higher complexity. Ecological models show that complexity usually destabilizes food webs (May 1973). This result empirically further corroborates the hypothesis that stability decreases with increasing number of trophic levels and omnivory (Borrelli and Ginzburg 2014). Food webs with many generalist species, like Beagle Channel’s, are less resistant to disturbances because, when affected, they tend to cause secondary extinctions (Wootton 2015). That is because generalists have many weak interactions which are known to be important for stability (McCann et al., 1998).

Although omnivory is not a direct measure of stability, it acts as a buffer to changes as the ecosystem presents alternative energy pathways in the face of perturbations. Omnivores are species able to adapt faster and at a wider range of environmental conditions by changing their foraging habits to feed on the most abundant prey (Fagan 1997). In this sense, the highly significant omnivore proportion in Beagle Channel food web suggests that the network could be more robust to variations in prey abundances than Potter Cove. Food web studies have found that the effect of omnivory on stability and measures of local stability (such as QSS) is influenced by species interaction’s strengths (Gellner and McCann 2012). Therefore, a thorough assessment of this effect would require knowledge on the distribution of interactions’ strength.

Modularity was different between the analyzed food webs, and even different compartmentalization mechanisms appear to be responsible in determining each food web structure. Compartmentalization has been proposed to arise as a result and combination of habitat heterogeneity within the environment, species functional group and the dependence of each node on energy derived from basal resources (Krause et al. 2003; Rezende et al. 2009; Zhao et al. 2018). Potter Cove had each of the four modules associated to an independent type of basal resource (1-macroalgae, 2-aged detritus, 3-fresh detritus/necromass/diatoms and 4-phytoplankton) with different habitats. These results support the important role energy pathways and habitat type play in structuring Antarctic food webs (Cordone et al. 2020). This pattern was not evident for Beagle Channel. Interestingly, the latter had one module with no primary productivity resource, meaning that it depends energetically on other modules, and made up of all the top predators (fish). The higher modularity of the sub-Antarctic food web is probably associated with a more heterogeneous and complex habitat created by the kelp forest (Teagle et al. 2017). This macroalgae species has strong non-trophic interactions (mutualism, competition) with others species (invertebrates, fish, other macroalgae), since it is not as important food resource as refugee and habitat (Miller et al., 2018; Riccialdelli et al., 2017), with indirect consequences for predator-prey interactions (Kéfi et al. 2012). To fully understand the processes shaping the structure of this sub-Antarctic ecosystem for future works we suggest the incorporation of non-trophic interactions into the network.

### 4.2 Food web similarities

We would have expected the different biogeographical, climatic and evolutionary history between Potter Cove and the Beagle Channel to be reflected in a marked different food web structure, although we found several similarities. The studied food webs showed the same connectance value (0.05) and similar proportions of basal, intermediate and top species. Marine food webs tend to resemble each other (Dunne, Williams, and Martinez 2004), when compared to non-marine ecosystem; and in high latitudes, they usually exhibit low connectance values (between 0.01 and 0.05) (De Santana et al. 2013; Kortsch et al. 2015). In this sense, our results support previous studies of polar food webs complexity.

Connectance usually covaries with ecosystem primary productivity and the proportion of basal species (Vermaat, Dunne, and Gilbert 2009). The similarities in connectance value, related to the presence of a very similar proportion of basal species, can be further explained by the importance of macroalgae as an energy source in both ecosystems. Macroalgae provide a direct pathway of energy and matter into organisms that feed on them and indirectly through the detritus pathway (Momo et al., 2020; Riccialdelli et al., 2017).

We found that Potter Cove and Beagle Channel food webs are in addition similarly compartmentalized in terms of number of modules. In both cases, the distribution of species topological roles showed that there is only one species responsible for linking modules and connecting the entire food web. However, such species have very different identities and trophic positions in each ecosystem. In Potter Cove the connector is the black rockcod (*Notothenia coriiceps)*, a demersal, generalist, omnivore and top predator fish (Barrera-Oro et al. 2019; Zamzow et al. 2011). For Beagle Channel it is the squat lobster (*Munida gregaria)*, a generalist in terms of habitat and prey items (Pérez-Barros et al. 2010; Vinuesa and Varisco 2007; Romero et al. 2004). This species has been already proposed as a wasp-waist species (a species with an intermediate trophic level that has high ecological importance by regulating the energy transfer between bottom and top trophic levels) for the Beagle Channel and adjacent areas (Riccialdelli et al., 2020). By feeding across many trophic levels and, in the squat lobster case, across pelagic and benthic habitats these species have a strong effect on food web connectivity, modularity and, therefore, stability. Any disturbance affecting these species could have catastrophic effects on the food web structure.

### 4.3 The impact of invasive species on food web structure

Invading species can alter the functioning of entire ecosystems (Convey and Peck 2019; Ehrenfeld 2010; Anderson and Rosemond 2010). Although great attention is being given to the prevention of biological invasions to Antarctica via the management and regulation of introduction pathways (Chown and Brooks 2019), species might still shift their distribution pushed by climate change, or arrive transported by human activities and invade new habitats (Hughes et al. 2020). Even more, it has been proposed that ecosystems with low connectance food webs (C∼0.05), like Potter Cove and Beagle Channel, are very vulnerable to invasions (Romanuk et al. 2009; Smith-Ramesh, Moore, and Schmitz 2017). The presence of exotic generalist species, many of which inhabit the Beagle Channel, was already reported for the WAP. One particular worrying case is the discovery of sub-Antarctic king crabs (Lithodidae) on the continental slope of the Antarctic Peninsula (Thatje and Arntz 2004). It is considered that the contemporary marine biota of Antarctica has been strongly shaped by the absence of durophagous (skeleton-crushing) predators for millions of years (Aronson et al. 2007). Viable populations of king crab generalist and durophagous predators (Lovrich & Vinuesa, 2016), would radically alter the structure of marine Antarctic food webs. It would increase connectance, since these species interact with multiple prey over space and time, while reducing food web compartmentalization (modularity) and, as a result, ecosystem resilience to perturbations (Stouffer and Bascompte 2011). In addition, recently a settlement of the bivalve *Mytilus sp*. was reported in the South Shetland Islands (Cárdenas et al. 2020), evidencing an invasion successful enough to allow for the development of the species under Antarctic temperatures. This mussel is known to be a remarkable invader and strong competitor, that can displace native sessile species, reducing biodiversity (Branch & Steffani, 2004; Thomsen et al., 2014), food web complexity and structure. In the Beagle Channel, the invasion of the chinook salmon could change the entire ecosystem structure. This species by potentially preying on top predator fish (Fernández et al. 2010) and the squat lobster (James and Unwin 1996) could cause a rapid secondary extinction cascade (Donohue et al. 2017).

## 5 Concluding remarks

The Beagle Channel food web can be summarized as more complex than Potter Cove’s, but less locally stable and sensitive to the loss of its most connected and generalist species. However, the high degree of omnivory and its stronger modularity suggest plasticity to adapt to changes before collapsing. The results for Potter Cove suggest a less complex structure, while presenting higher probability to recover after a perturbation. Despite showing different architecture, our result suggests that the Antarctic and sub-Antarctic food webs evolved so that different structural characteristics can provide mechanisms to cope in the face of perturbations.

In a time of rapid anthropogenic and climate changes, that causes extensive ecosystem transformations, it is crucial to explore trophic interactions in a network framework, and understand their influence in ecosystem functioning. This study highlights the powerful tool network analysis applied to food webs, since it has allowed us to identify similarities and differences among two seemingly different complex systems. Our results provide a baseline of information on the comparison of food web structure and stability for Antarctic and sub-Antarctic ecosystems, that could be used as a management tool to evaluate future modifications resulting from biological invasions and other anthropogenic impacts and climate change.

## 7 Data Availability Statement

All source code and data that support the findings of this study are openly available in GitHub at https://github.com/123iamela/foodweb-comparison and Zenodo at doi:10.5281/zenodo.4688715.

## 8 Author Contributions

I.D.R, I.R.S and L.A.S. conceived the study. I.D.R and T.I.M. compiled the data. I.D.R and L.A.S designed and performed the analytical calculations and produced the figures. I.D.R. wrote the initial draft of the manuscript. T.I.M, I.R.S and L.A.S. reviewed, edited, and revised the manuscript. All authors contributed to the manuscript and approved the submitted version.

## 10 Acknowledgements

The manuscript was submitted as a preprint at bioRxiv, doi: 10.1101/2021.04.12.439560.

## References

Adami, Mariana L, and Sandra Gordillo. 1999. “Structure and Dynamics of the Biota Associated with Macrocystis Pyrifera (Phaeophyta) from the Beagle Channel, Tierra Del Fuego.” Scientia Marina 63 (1889): 183–91. https://doi.org/10.3989/scimar.1999.63s1183.

Aguirre, Gastón E., Fabiana L. Capitanio, Gustavo A. Lovrich, and Graciela B. Esnal. 2012. “Seasonal Variability of Metazooplankton in Coastal Sub-Antarctic Waters (Beagle Channel).” Marine Biology Research 8 (4): 341–53. https://doi.org/10.1080/17451000.2011.627922.

Albert, Réka, Hawoong Jeong, and Albert László Barabási. 2000. “Error and Attack Tolerance of Complex Networks.” Nature 406 (6794): 378–82. https://doi.org/10.1038/35019019.

Allesina, Stefano, and Mercedes Pascual. 2008. “Network Structure, Predator - Prey Modules, and Stability in Large Food Webs.” Theoretical Ecology 1 (1): 55–64. https://doi.org/10.1007/s12080-007-0007-8.

Almandoz, Gastón O., Marcelo P. Hernando, Gustavo A. Ferreyra, Irene R. Schloss, and Martha E. Ferrario. 2011. “Seasonal Phytoplankton Dynamics in Extreme Southern South America (Beagle Channel, Argentina).” Journal of Sea Research 66 (2): 47–57. https://doi.org/10.1016/j.seares.2011.03.005.

Amin, Oscar, Laura Comoglio, Carla Spetter, Claudia Duarte, Raúl Asteasuain, Rubén Hugo Freije, and Jorge Marcovecchio. 2011. “Assessment of Land Influence on a High-Latitude Marine Coastal System: Tierra Del Fuego, Southernmost Argentina.” Environmental Monitoring and Assessment 175 (1–4): 63–73. https://doi.org/10.1007/s10661-010-1493-5.

Amin, Oscar, Laura Ferrer, and Jorge Marcovecchio. 1996. “Heavy Metal Concentrations in Litteral Sediments from the Beagle Channel, Tierra Del Fuego, Argentina.” Environmental Monitoring and Assessment 41 (3): 219–31. https://doi.org/10.1007/BF00419743.

Anderson, Christopher B., and Amy D. Rosemond. 2010. “Beaver Invasion Alters Terrestrial Subsidies to Subantarctic Stream Food Webs.” Hydrobiologia 652 (1): 349–61. https://doi.org/10.1007/s10750-010-0367-8.

Arntz, Wolf E., and Matthias Gorny. 1994. “Cruise Report of the Joint Chilean-German-Italian Magellan ‘Victor Hensen’ Campaign in 1994.” Ber.Polarforsch. 190 (1996): 1–116. https://doi.org/10.2312/BzP_0190_1996.

Aronson, Richard B., Sven Thatje, Andrew Clarke, Lloyd S. Peck, Daniel B. Blake, Cheryl D. Wilga, and Brad A. Seibel. 2007. “Climate Change and Invasibility of the Antarctic Benthos.” Annual Review of Ecology, Evolution, and Systematics 38 (1): 129–54. https://doi.org/10.1146/annurev.ecolsys.38.091206.095525.

Balestrini, C, G Manzella, and G Lovrich. 1998. “Simulación de Corrientes En El Canal Beagle y Bahia Ushuaia, Mediante Un Modelo Bidimensional.” Servicio de Hidrografia Naval 98 (January): 1–58. https://doi.org/10.13140/RG.2.1.1196.2729.

Barker, P. F., and E. Thomas. 2004. “Origin, Signature and Palaeoclimatic Influence of the Antarctic Circumpolar Current.” Earth-Science Reviews 66 (1–2): 143–62. https://doi.org/10.1016/j.earscirev.2003.10.003.

Barrera-Oro, Esteban. 2002. “The Role of Fish in the Antarctic Marine Food Web: Differences between Inshore and Offshore Waters in the Southern Scotia Arc and West Antarctic Peninsula.” Antarctic Science 14 (4): 293–309. https://doi.org/10.1017/S0954102002000111.

Barrera-Oro, Esteban, Eugenia Moreira, Meike Anna Seefeldt, Mariano Valli Francione, and María Liliana Quartino. 2019. “The Importance of Macroalgae and Associated Amphipods in the Selective Benthic Feeding of Sister Rockcod Species Notothenia Rossii and N. Coriiceps (Nototheniidae) in West Antarctica.” Polar Biology 42 (2): 317–34. https://doi.org/10.1007/s00300-018-2424-0.

Barrera Oro Esteban, and Ricardo Jorge Casaux. 2008. “General Ecology of Coastal Fish from the South Shetland Island and West Antarctic Peninsula Areas.” In The Antarctic Ecosystem of Potter Cove, King-George Island (Isla 25 de Mayo), edited by C. Wiencke, G.A. Ferreyra, D. Abele, and S. Marenssi, 95–110. Alfred Wegener Institut für Polar und Meeresforschung.

Blois, Jessica L., Phoebe L. Zarnetske, Matthew C. Fitzpatrick, and Seth Finnegan. 2013. “Climate Change and the Past, Present, and Future of Biotic Interactions.” Science 341 (6145): 499–504. https://doi.org/10.1126/science.1237184.

Borrelli, Jonathan J., and Lev R. Ginzburg. 2014. “Why There Are so Few Trophic Levels: Selection against Instability Explains the Pattern.” Food Webs 1 (1–4): 10–17. https://doi.org/10.1016/j.fooweb.2014.11.002.

Branch, George M., and C. Nina Steffani. 2004. “Can We Predict the Effects of Alien Species? A Case-History of the Invasion of South Africa by Mytilus Galloprovincialis (Lamarck).” Journal of Experimental Marine Biology and Ecology 300 (1–2): 189–215. https://doi.org/10.1016/j.jembe.2003.12.007.

Briand, Frédéric, and Joel E. Cohen. 1984. “Community Food Webs Have Scale-Invariant Structure.” Nature 307 (5948): 264–67. https://doi.org/10.1038/307264a0.

Bruno, Daniel O., Mariela F. Victorio, Eduardo M. Acha, and Daniel A. Fernández. 2018. “Fish Early Life Stages Associated with Giant Kelp Forests in Sub-Antarctic Coastal Waters (Beagle Channel, Argentina).” Polar Biology 41 (2): 365–75. https://doi.org/10.1007/s00300-017-2196-y.

Burnham, Kenneth P., and David R. Anderson. 2002. Model Selection and Multimodel Inference: A Practical Information-Theoritic Approach. Second Edition. Springer. https://doi.org/10.1007/978-0-387-22456-5_7.

Cárdenas, Leyla, Jean Charles Leclerc, Paulina Bruning, Ignacio Garrido, Camille Détrée, Alvaro Figueroa, Marcela Astorga, et al. 2020. “First Mussel Settlement Observed in Antarctica Reveals the Potential for Future Invasions.” Scientific Reports 10 (1): 1–8. https://doi.org/10.1038/s41598-020-62340-0.

Castilla, J. C. 1985. “Food Webs and Functional Aspects of the Kelp, Macrocystis Pyrifera, Community in the Beagle Channel, Chile.” Antarctic Nutrient Cycles and Food Webs. https://doi.org/10.1007/978-3-642-82275-9_57.

Chown, Steven L., and Cassandra M. Brooks. 2019. “The State and Future of Antarctic Environments in a Global Context.” Annual Review of Environment and Resources 44: 1–30. https://doi.org/10.1146/annurev-environ-101718-033236.

Clarke, Andrew, and Colin M. Harris. 2003. “Polar Marine Ecosystems: Major Threats and Future Change.” Environmental Conservation 30 (1): 1–25. https://doi.org/10.1017/S0376892903000018.

Convey, Peter, and Lloyd S. Peck. 2019. “Antarctic Environmental Change and Biological Responses.” Science Advances 5 (11). https://doi.org/10.1126/sciadv.aaz0888.

Cook, A. J., A. J. Fox, D. G. Vaughan, and J. G. Ferrigno. 2005. “Retreating Glacier Fronts on the Antarctic Peninsula over the Past Half-Century.” Science 308 (5721): 541–44. https://doi.org/10.1126/science.1104235.

Cordone, Georgina, Tomás I. Marina, Vanesa Salinas, Santiago R. Doyle, Leonardo A. Saravia, and Fernando R. Momo. 2018. “Effects of Macroalgae Loss in an Antarctic Marine Food Web: Applying Extinction Thresholds to Food Web Studies.” PeerJ 6: e5531. https://doi.org/10.7717/peerj.5531.

Cordone, Georgina, Vanesa Salinas, Tomás I. Marina, Santiago R. Doyle, Francesca Pasotti, Leonardo A. Saravia, and Fernando R. Momo. 2020. “Green vs Brown Food Web: Effects of Habitat Type on Multidimensional Stability Proxies for a Highly-Resolved Antarctic Food Web.” Food Webs, September, e00166. https://doi.org/10.1016/j.fooweb.2020.e00166.

Delmas, Eva, Mathilde Besson, Marie-Helene Brice, Laura Burkle, Giulio V Dalla Riva, Marie-Josee Fortin, Dominique Gravel, et al. 2018. “Analyzing Ecological Networks of Species Interactions.” BioRxiv, 112540. https://doi.org/10.1101/112540.

Díaz Claudia Andrade. 2016. “Trophic Structures and Flows in Marine Benthic Communities of the Magellan Region, Southern Chile,” 1–104.

Diez, Mariano J., and Gustavo A. Lovrich. 2010. “Reproductive Biology of the Crab Halicarcinus Planatus (Brachyura, Hymenosomatidae) in Sub-Antarctic Waters.” Polar Biology 33 (3): 389–401. https://doi.org/10.1007/s00300-009-0716-0.

Donohue, Ian, Owen L. Petchey, José M. Montoya, Andrew L. Jackson, Luke Mcnally, Mafalda Viana, Kevin Healy, Miguel Lurgi, Nessa E. O’Connor, and Mark C. Emmerson. 2013. “On the Dimensionality of Ecological Stability.” Ecology Letters 16 (4): 421–29. https://doi.org/10.1111/ele.12086.

Donohue, Ian, Owen L Petchey, Sonia Kéfi, Alexandre Génin, Andrew L Jackson, Qiang Yang, and Nessa E O’Connor. 2017. “Loss of Predator Species, Not Intermediate Consumers, Triggers Rapid and Dramatic Extinction Cascades.” Global Change Biology 0 (March): 1–11. https://doi.org/10.1111/gcb.13703.

Ducklow, Hugh W., William R. Fraser, Michael P. Meredith, Sharon E. Stammerjohn, Scott C. Doney, Douglas G. Martinson, Sévrine F. Sailley, et al. 2013. “West Antarctic Peninsula: An Ice-Dependent Coastal Marine Ecosystem in Transition.” Oceanography 26 (3): 190–203. https://doi.org/10.5670/oceanog.2013.62.

Duffy, J. Emmett, Casey M. Godwin, and Bradley J. Cardinale. 2017. “Biodiversity Effects in the Wild Are Common and as Strong as Key Drivers of Productivity.” Nature 549 (7671): 261–64. https://doi.org/10.1038/nature23886.

Dunne, Jennifer A. 2006. Ecological Networks: Linking Structure to Dynamics in Food Webs.

Dunne, Jennifer A., Richard J. Williams, and Neo D. Martinez. 2004. “Network Structure and Robustness of Marine Food Webs.” Marine Ecology Progress Series 273: 291–302. https://doi.org/10.3354/meps273291.

Dunne, Jennifer A, Richard J Williams, and Neo D Martinez. 2002. “Food-Web Structure and Network Theory: The Role of Connectance and Size.” Proceedings of the National Academy of Sciences 99 (20): 12917–22. https://doi.org/10.1073/pnas.192407699.

Ehrenfeld, Joan G. 2010. “Ecosystem Consequences of Biological Invasions.” Annual Review of Ecology, Evolution, and Systematics 41: 59–80. https://doi.org/10.1146/annurev-ecolsys-102209-144650.

Estrada, Ernesto. 2007. “Food Webs Robustness to Biodiversity Loss: The Roles of Connectance, Expansibility and Degree Distribution.” Journal of Theoretical Biology 244 (2): 296–307. https://doi.org/10.1016/j.jtbi.2006.08.002.

Fagan, William F. 1997. “Omnivory as a Stabilizing Feature of Natural Communities.” American Naturalist 150 (5): 554–67. https://doi.org/10.1086/286081.

Fernández Daniel Alfredo, Javier Ciancio, Santiago Guillermo Ceballos, Carla Riva-Rossi, and Miguel Alberto Pascual. 2010. “Chinook Salmon (Oncorhynchus Tshawytscha, Walbaum 1792) in the Beagle Channel, Tierra Del Fuego: The Onset of an Invasion.” Biological Invasions 12 (9): 2991–97. https://doi.org/10.1007/s10530-010-9731-x.

Frank, Kenneth T., Brian Petrie, Jae S. Choi, and William C. Leggett. 2005. “Ecology: Trophic Cascades in a Formerly Cod-Dominated Ecosystem.” Science 308 (5728): 1621–23. https://doi.org/10.1126/science.1113075.

Fraser, Ceridwen I., Adele K. Morrison, Andrew Mc C. Hogg, Erasmo C. Macaya, Erik van Sebille, Peter G. Ryan, Amanda Padovan, Cameron Jack, Nelson Valdivia, and Jonathan M. Waters. 2018. “Antarctica’s Ecological Isolation Will Be Broken by Storm-Driven Dispersal and Warming.” Nature Climate Change 8 (8): 704–8. https://doi.org/10.1038/s41558-018-0209-7.

Fuentes, Verónica, Gastón Alurralde, Bettina Meyer, Gastón E. Aguirre, Antonio Canepa, Anne Cathrin Wölfl, H. Christian Hass, Gabriela N. Williams, and Irene R. Schloss. 2016. “Glacial Melting: An Overlooked Threat to Antarctic Krill.” Scientific Reports 6 (June): 1–12. https://doi.org/10.1038/srep27234.

Gellner, Gabriel, and Kevin McCann. 2012. “Reconciling the Omnivory-Stability Debate.” American Naturalist 179 (1): 22–37. https://doi.org/10.1086/663191.

Gibert, Jean P. 2019. “Temperature Directly and Indirectly Influences Food Web Structure.” Scientific Reports 9 (1): 1–8. https://doi.org/10.1038/s41598-019-41783-0.

Gil, M N, A I Torres, O Amin, and J L Esteves. 2011. “Assessment of Recent Sediment Influence in an Urban Polluted Subantarctic Coastal Ecosystem. Beagle Channel (Southern Argentina).” Marine Pollution Bulletin 62 (1): 201–7. https://doi.org/10.1016/j.marpolbul.2010.10.004.

Gillespie, Colin S. 2015. “Fitting Heavy Tailed Distributions: The Powerlaw Package.” Journal of Statistical Software. https://doi.org/10.18637/jss.v064.i02.

Graham, Michael H, Julio A Vásquez, and Alejandro H Buschmann. 2007. “Global Ecology of the Giant Kelp Macrocystis: From Ecotypes to Ecosystems.” Oceanography and Marine Biology 45: 39–88.

Griffiths, Huw James, Andrew J S Meijers, and Thomas J Bracegirdle. 2017. “More Losers than Winners in a Century of Future.” Nature Climate Change 7 (September).

Griffiths, Huw James, and Catherine Louise Waller. 2016. “The First Comprehensive Description of the Biodiversity and Biogeography of Antarctic and Sub-Antarctic Intertidal Communities.” Journal of Biogeography 43 (6): 1143–55. https://doi.org/10.1111/jbi.12708.

Grilli, Jacopo, Tim Rogers, and Stefano Allesina. 2016. “Modularity and Stability in Ecological Communities.” Nature Communications 7 (May): 1–10. https://doi.org/10.1038/ncomms12031.

Guimera, Roger. 2005. “Functional Cartography of Complex Metabolic Networks” 433 (February): 895–900. https://doi.org/10.1038/nature03286.1.

Gutt, Julian, Nancy Bertler, Thomas J. Bracegirdle, Alexander Buschmann, Josefino Comiso, Graham Hosie, Enrique Isla, et al. 2015. “The Southern Ocean Ecosystem under Multiple Climate Change Stresses - an Integrated Circumpolar Assessment.” Global Change Biology 21 (4): 1434–53. https://doi.org/10.1111/gcb.12794.

Hickling, Rachael, David B. Roy, Jane K. Hill, Richard Fox, and Chris D. Thomas. 2006. “The Distributions of a Wide Range of Taxonomic Groups Are Expanding Polewards.” Global Change Biology 12 (3): 450–55. https://doi.org/10.1111/j.1365-2486.2006.01116.x.

Hoegh-Guldberg, Ove, and John F. Bruno. 2010. “The Impact of Climate Change On.” Science 328 (June): 1523–29. https://doi.org/10.1367/1539-4409(2003)003<0044:TIOCCO>2.0.CO;2.

Hughes, Kevin A., Oliver L. Pescott, Jodey Peyton, Tim Adriaens, Elizabeth J. Cottier-Cook, Gillian Key, Wolfgang Rabitsch, et al. 2020. “Invasive Non-Native Species Likely to Threaten Biodiversity and Ecosystems in the Antarctic Peninsula Region.” Global Change Biology 26 (4): 2702–16. https://doi.org/10.1111/gcb.14938.

IPCC Working Group 1, I. T.F. Stocker, D. Qin, G.-K. Plattner, M. Tignor, S.K. Allen, J. Boschung, et al. 2013. “IPCC, 2013: Climate Change 2013: The Physical Science Basis. Contribution of Working Group I to the Fifth Assessment Report of the Intergovernmental Panel on Climate Change.” IPCC.

James, Gavin D., and Martin J. Unwin. 1996. “Diet of Chinook Salmon (Oncorhynchus Tshawytscha) in Canterbury Coastal Waters, New Zealand.” New Zealand Journal of Marine and Freshwater Research 30 (1): 69–78. https://doi.org/10.1080/00288330.1996.9516697.

Kéfi, Sonia, Eric L. Berlow, Evie A. Wieters, Sergio A. Navarrete, Owen L. Petchey, Spencer A. Wood, Alice Boit, et al. 2012. “More than a Meal… Integrating Non-Feeding Interactions into Food Webs.” Ecology Letters 15 (4): 291–300. https://doi.org/10.1111/j.1461-0248.2011.01732.x.

Kéfi, Sonia, Vincent Miele, Evie A. Wieters, Sergio A. Navarrete, and Eric L. Berlow. 2016. “How Structured Is the Entangled Bank? The Surprisingly Simple Organization of Multiplex Ecological Networks Leads to Increased Persistence and Resilience.” PLoS Biology 14 (8): 1–21. https://doi.org/10.1371/journal.pbio.1002527.

Kortsch, Susanne, Raul Primicerio, Michaela Aschan, Sigrid Lind, Andrey V. Dolgov, and Benjamin Planque. 2018. “Food-Web Structure Varies along Environmental Gradients in a High-Latitude Marine Ecosystem.” Ecography, 1–14. https://doi.org/10.1111/ecog.03443.

Kortsch, Susanne, Raul Primicerio, Maria Fossheim, Andrey V. Dolgov, and Michaela Aschan. 2015. “Climate Change Alters the Structure of Arctic Marine Food Webs Due to Poleward Shifts of Boreal Generalists.” Proceedings of the Royal Society B: Biological Sciences 282 (1814): 20151546. https://doi.org/10.1098/rspb.2015.1546.

Krause, Ann E, Kenneth A Frank, Doran M Mason, Robert E Ulanowicz, and William W Taylor. 2003. “Compartments Revealed in Food-Web Structure” 426 (November).

Lagger, Cristian, Mónica Nime, Luciana Torre, Natalia Servetto, Marcos Tatián, and Ricardo Sahade. 2017. “Climate Change, Glacier Retreat and a New Ice-Free Island Offer New Insights on Antarctic Benthic Responses.” Ecography, n/a--n/a. https://doi.org/10.1111/ecog.03018.

Landi, Pietro, Henintsoa O. Minoarivelo, Å ke Brännström, Cang Hui, and Ulf Dieckmann. 2018. “Complexity and Stability of Ecological Networks: A Review of the Theory.” Population Ecology 60 (4): 319–45. https://doi.org/10.1007/s10144-018-0628-3.

López-Farrán, Zambra, Charlène Guillaumot, Luis Vargas-Chacoff, Kurt Paschke, Valérie Dulière, Bruno Danis, Elie Poulin, Thomas Saucède, Jonathan Waters, and Karin Gérard. 2021. “Is the Southern Crab Halicarcinus Planatus (Fabricius, 1775) the next Invader of Antarctica?” Global Change Biology. https://doi.org/10.1111/gcb.15674.

Lovrich, Gustavo A. 1997. “La Pesquería Mixta de Las Centollas Lithodes Santolla y Paralomis Granulosa (Anomura: Lithodidae) En Tierra Del Fuego, Argentina.” Investigaciones Marinas 25: 41–57. https://doi.org/10.4067/S0717-71781997002500004.

Lovrich, Gustavo Alejandro, and Julio H Vinuesa. 2016. “BIOLOGÍ A DE LAS CENTOLLAS (ANOMURA : LITHODIDAE).” EL MAR ARGENTINO Y SUS RECURSOS PESQUEROS 6 (October): 183–212.

Marina, Tomás I., Vanesa Salinas, Georgina Cordone, Gabriela Campana, Eugenia Moreira, Dolores Deregibus, Luciana Torre, et al. 2018. “The Food Web of Potter Cove (Antarctica): Complexity, Structure and Function.” Estuarine, Coastal and Shelf Science 200: 141–51. https://doi.org/10.1016/j.ecss.2017.10.015.

Massey, Frank Jr. 1951. “The Kolmogorov-Smirnov Test for Goodness of Fit.” Journal of the American Statistical Association.

May, R. M. 1973. “Stability and Complexity in Model Ecosystems.” Monographs in Population Biology. https://doi.org/10.2307/3743.

Mccallum, Hamish. 2008. Population Parameters: Estimation for Ecological Models. Population Parameters: Estimation for Ecological Models. https://doi.org/10.1002/9780470757468.

McCann, K., and A. Hastings. 1997. “Re-Evaluating the Omnivory-Stability Relationship in Food Webs.” Proceedings of the Royal Society B: Biological Sciences 264 (1385): 1249–54. https://doi.org/10.1098/rspb.1997.0172.

McCann, Kevin, Alan Hastings, and Gary R. Huxel. 1998. “Weak Trophic Interactions and the Balance of Nature.” Nature 395 (6704): 794–98. https://doi.org/10.1038/27427.

McCarthy, Arlie H., Lloyd S. Peck, Kevin A. Hughes, and David C. Aldridge. 2019. “Antarctica: The Final Frontier for Marine Biological Invasions.” Global Change Biology 25 (7): 2221–41. https://doi.org/10.1111/gcb.14600.

Meredith, Michael P, M. Sommerkorn, S. Cassotta, C. Derksen, A. Ekaykin, A. Hollowed, G. Kofinas, et al. 2019. “Polar Regions. In: IPCC Special Report on the Ocean and Cryosphere in a Changing Climate.” IPCC. https://www.ipcc.ch/srocc/chapter/chapter-3-2/.

Miller, Robert J., Kevin D. Lafferty, Thomas Lamy, Li Kui, Andrew Rassweiler, and Daniel C. Reed. 2018. “Giant Kelp, Macrocystis Pyrifera, Increases Faunal Diversity through Physical Engineering.” Proceedings of the Royal Society B: Biological Sciences 285 (1874). https://doi.org/10.1098/rspb.2017.2571.

Momo, Fernando R, Georgina Cordone, Tomás I Marina, Vanesa Salinas, Gabriela L Campana, Mariano A Valli, Santiago R Doyle, and Leonardo A Saravia. 2020. “Seaweeds in the Antarctic Marine Coastal Food Web.” In Antarctic Seaweeds: Diversity, Adaptation and Ecosystem Services, edited by Iván Gómez and Pirjo Huovinen, 293–307. Cham: Springer International Publishing. https://doi.org/10.1007/978-3-030-39448-6_15.

Montes-Hugo, Martin, Scott C. Doney, Hugh W. Ducklow, William R. Fraser, Douglas Martinson, Sharon E. Stammerjohn, and Oscar Schofield. 2009. “Recent Changes in Phytoplankton Communities Associated with Rapid Regional Climate Change along the WAP” 323 (March).

Montoya, José M., Stuart L. Pimm, and Ricard V. Solé. 2006. “Ecological Networks and Their Fragility.” Nature 442 (7100): 259–64. https://doi.org/10.1038/nature04927.

Mora, Bernat Bramon, Dominique Gravel, Luis J. Gilarranz, Timothée Poisot, and Daniel B. Stouffer. 2018. “Identifying a Common Backbone of Interactions Underlying Food Webs from Different Ecosystems.” Nature Communications 9 (1). https://doi.org/10.1038/s41467-018-05056-0.

Moreno, CA, and HF Jara. 1984. “Ecological Studies on Fish Fauna Associated with Macrocystis Pyrifera Belts in the South of Fueguian Islands, Chile.” Marine Ecology Progress Series 15 (October 1984): 99–107. https://doi.org/10.3354/meps015099.

Nardi, Cristina Fernanda, Daniel Alfredo Fernández, Fabián Alberto Vanella, and Tomás Chalde. 2019. “The Expansion of Exotic Chinook Salmon (Oncorhynchus Tshawytscha) in the Extreme South of Patagonia: An Environmental DNA Approach.” Biological Invasions 21 (4): 1415–25. https://doi.org/10.1007/s10530-018-1908-8.

Pasotti, Francesca, Elena Manini, Donato Giovannelli, Anne Cathrin Wölfl, Donata Monien, Elie Verleyen, Ulrike Braeckman, Doris Abele, and Ann Vanreusel. 2015. “Antarctic Shallow Water Benthos in an Area of Recent Rapid Glacier Retreat.” Marine Ecology 36 (3): 716–33. https://doi.org/10.1111/maec.12179.

Pérez-Barros, Patricia, M. Carolina Romero, Javier A Calcagno, and Gustavo A Lovrich. 2010. “Similar Feeding Habits of Two Morphs of Munida Gregaria (Decapoda) Evidence the Lack of Trophic Polymorphism.” Revista de Biología Marina y Oceanografía 45 (3): 461–70. https://doi.org/10.4067/s0718-19572010000300011.

R Development Core Team, R. 2020. R: A Language and Environment for Statistical Computing. R Foundation for Statistical Computing. https://doi.org/10.1007/978-3-540-74686-7.

Rezende, Enrico L., Eva M. Albert, Miguel A. Fortuna, and Jordi Bascompte. 2009. “Compartments in a Marine Food Web Associated with Phylogeny, Body Mass, and Habitat Structure.” Ecology Letters 12 (8): 779–88. https://doi.org/10.1111/j.1461-0248.2009.01327.x.

Riccialdelli, L, YA Becker, NE Fioramonti, M Torres, DO Bruno, A Raya Rey, and DA Fernández. 2020. “Trophic Structure of Southern Marine Ecosystems: A Comparative Isotopic Analysis from the Beagle Channel to the Oceanic Burdwood Bank Area under a Wasp-Waist Assumption.” Marine Ecology Progress Series 655: 1–27. https://doi.org/10.3354/meps13524.

Riccialdelli, Luciana, Seth D. Newsome, Marilyn L. Fogel, and Daniel A. Fernández. 2017. “Trophic Interactions and Food Web Structure of a Subantarctic Marine Food Web in the Beagle Channel: Bahía Lapataia, Argentina.” Polar Biology 40 (4): 807–21. https://doi.org/10.1007/s00300-016-2007-x.

Rodriguez, Iara Diamela, Tomás Ignacio Marina, Irene Ruth Schloss, and Leonardo Ariel Saravia. 2021. “Marine Food Webs Are More Complex but Less Stable in Sub-Antarctic than in Antarctic Regions.” BioRxiv. https://doi.org/10.1101/2021.04.12.439560.

Romanuk, Tamara N., Yun Zhou, Ulrich Brose, Eric L. Berlow, Richard J. Williams, and Neo D. Martinez. 2009. “Predicting Invasion Success in Complex Ecological Networks.” Philosophical Transactions of the Royal Society B: Biological Sciences 364 (1524): 1743–54. https://doi.org/10.1098/rstb.2008.0286.

Romero, M. Carolina, Gustavo A. Lovrich, Federico Tapella, and Sven Thatje. 2004. “Feeding Ecology of the Crab Munida Subrugosa (Decapoda: Anomura: Galatheidae) in the Beagle Channel, Argentina.” Journal of the Marine Biological Association of the United Kingdom 84 (2): 359–65. https://doi.org/10.1017/S0025315404009282h.

Sahade, Ricardo, Cristian Lagger, Luciana Torre, Fernando Momo, Patrick Monien, Irene Schloss, David K.A. A Barnes, et al. 2015. “Climate Change and Glacier Retreat Drive Shifts in an Antarctic Benthic Ecosystem.” Science Advances 1 (10): e1500050. https://doi.org/10.1126/sciadv.1500050.

Saint-Béat, Blanche, Dan Baird, Harald Asmus, Ragnhild Asmus, Cédric Bacher, Stephen R. Pacella, Galen A. Johnson, Valérie David, Alain F. Vézina, and Nathalie Niquil. 2015. “Trophic Networks: How Do Theories Link Ecosystem Structure and Functioning to Stability Properties? A Review.” Ecological Indicators 52: 458–71. https://doi.org/10.1016/j.ecolind.2014.12.017.

Santana, Charles N. De, Alejandro F. Rozenfeld, Pablo A. Marquet, and Carlos M. Duart. 2013. “Topological Properties of Polar Food Webs.” Marine Ecology Progress Series 474: 15–26. https://doi.org/10.3354/meps10073.

Saravia, Leonardo A., Susanne Kortsch, Tomás I. Marina, and Iara D. Rodriguez. 2019. “Multiweb: R Package for Multiple Interaction Ecological Networks.” https://doi.org/10.5281/ZENODO.3370397.

Schloss, Irene R., Gustavo A. Ferreyra, and Diana Ruiz-Pino. 2002. “Phytoplankton Biomass in Antarctic Shelf Zones: A Conceptual Model Based on Potter Cove, King George Island.” Journal of Marine Systems 36 (3–4): 129–43. https://doi.org/10.1016/S0924-7963(02)00183-5.

Schloss, Irene R, Doris Abele, Sébastien Moreau, Serge Demers, A Valeria Bers, Oscar González, and Gustavo A Ferreyra. 2012. “Response of Phytoplankton Dynamics to 19-Year (1991-2009) Climate Trends in Potter Cove (Antarctica).” Journal of Marine Systems 92 (1): 53–66. https://doi.org/10.1016/j.jmarsys.2011.10.006.

Smith-Ramesh, Lauren M., Alexandria C. Moore, and Oswald J. Schmitz. 2017. “Global Synthesis Suggests That Food Web Connectance Correlates to Invasion Resistance.” Global Change Biology 23 (2): 465–73. https://doi.org/10.1111/gcb.13460.

Song, Chuliang, and Serguei Saavedra. 2020. “Telling Ecological Networks Apart by Their Structure: An Environment-Dependent Approach.” PLoS Computational Biology 16 (4): 1–15. https://doi.org/10.1371/journal.pcbi.1007787.

Steig, Eric J., David P. Schneider, Scott D. Rutherford, Michael E. Mann, Josefino C. Comiso, and Drew T. Shindell. 2009. “Warming of the Antarctic Ice-Sheet Surface since the 1957 International Geophysical Year.” Nature 457 (7228): 459–62. https://doi.org/10.1038/nature07669.

Stouffer, Daniel B., and Jordi Bascompte. 2011. “Compartmentalization Increases Food-Web Persistence.” Proceedings of the National Academy of Sciences of the United States of America 108 (9): 3648–52. https://doi.org/10.1073/pnas.1014353108.

Strona, Giovanni, Domenico Nappo, Francesco Boccacci, Simone Fattorini, and Jesus San-Miguel-Ayanz. 2014. “A Fast and Unbiased Procedure to Randomize Ecological Binary Matrices with Fixed Row and Column Totals.” Nature Communications 5 (May): 1–7. https://doi.org/10.1038/ncomms5114.

Tapella, F., M.C. Romero, G.A. Lovrich, and A. Chizzini. 2002. “Life History of the Galatheid Crab Munida Subrugosa in Subantarctic Waters of the Beagle Channel, Argentina,” no. January 2014: 115–34. https://doi.org/10.4027/ccwrbme.2002.11.

Teagle, Harry, Stephen J. Hawkins, Pippa J. Moore, and Dan A. Smale. 2017. “The Role of Kelp Species as Biogenic Habitat Formers in Coastal Marine Ecosystems.” Journal of Experimental Marine Biology and Ecology 492: 81–98. https://doi.org/10.1016/j.jembe.2017.01.017.

Thatje, Sven, and Wolf E. Arntz. 2004. “Antarctic Reptant Decapods: More than a Myth?” Polar Biology 27 (4): 195–201. https://doi.org/10.1007/s00300-003-0583-z.

The International Association of Antarctica Tour Operators (IAATO). 2019. “Report Tourism in Antarctica 2019.” https://iaato.org/wp-content/uploads/2020/04/Tourism_in_Antarctica_2019.pdf.

Thompson, Ross M, Martin Hemberg, Brian M Starzomski, and Jonathan B Shurin. 2007. “Trophic Levels and Trophic Tangles: The Prevalence of Omnivory in Real Food Webs.” Ecology 88 (3): 612–17. http://www.ncbi.nlm.nih.gov/pubmed/17503589.

Thomsen, Mads S., James E. Byers, David R. Schiel, John F. Bruno, Julian D. Olden, Thomas Wernberg, and Brian R. Silliman. 2014. “Impacts of Marine Invaders on Biodiversity Depend on Trophic Position and Functional Similarity.” Marine Ecology Progress Series 495: 39–47. https://doi.org/10.3354/meps10566.

Tilman, David. 1996. “Biodiversity: Population Versus Ecosystem Stability.” Ecology 77 (2): 350–63. https://doi.org/10.2307/2265614.

Turner, John, Nicholas E. Barrand, Thomas J. Bracegirdle, Peter Convey, Dominic A. Hodgson, Martin Jarvis, Adrian Jenkins, et al. 2014. “Antarctic Climate Change and the Environment: An Update.” Polar Record 50 (3): 237–59. https://doi.org/10.1017/S0032247413000296.

Vermaat, Jan E., Jennifer A. Dunne, and Alison J. Gilbert. 2009. “Major Dimensions in Food-Web Structure Properties.” Ecology 90 (1): 278–82. https://doi.org/10.1890/07-0978.1.

Vinuesa, Julio H, and Martin Varisco. 2007. “Trophic Ecology of the Lobster Krill Munida Gregaria in San Jorge Gulf, Argentina.” Investigaciones Marinas 35 (2): 25–34. https://doi.org/10.4067/S0717-71782007000200003.

Wootton, K. L. 2015. “Fitting Species into the Complexity-Stability Debate.” University of Canterbury, New Zealand.

Wootton, K.L. 2017. “Omnivory and Stability in Freshwater Habitats: Does Theory Match Reality?” Freshwater Biology 62 (5): 821–32. https://doi.org/10.1111/fwb.12908.

Young, Hillary S, Douglas J Mccauley, Robert B Dunbar, Michael S Hutson, Miller Ter-kuile, Rodolfo Dirzo, S Young, J Mccauley, and S Hutson. 2013. “The Roles of Productivity and Ecosystem Size in Determining Food Chain Length in Tropical Terrestrial Ecosystems” 94 (3): 692–701.

Zamzow, Jill P., Craig F. Aumack, Charles D. Amsler, James B. McClintock, Margaret O. Amsler, and Bill J. Baker. 2011. “Gut Contents and Stable Isotope Analyses of the Antarctic Fish, Notothenia Coriiceps (Richardson), from Two Macroalgal Communities.” Antarctic Science 23 (2): 107–16. https://doi.org/10.1017/S095410201000091X.

Zhao, Lei, Huayong Zhang, Wang Tian, and Xiang Xu. 2018. “Identifying Compartments in Ecological Networks Based on Energy Channels.” Ecology and Evolution 8 (1): 309–18. https://doi.org/10.1002/ece3.3648.

