## Supplemental Information for "Marine food webs are more complex but less stable in sub-Antarctic than in Antarctic regions"

**Equation 1: Connectance (C)**

$C = \frac{L}{S^2}$ (Eq.1)

Where *L* is the number of trophic interactions and *S* the number of species.

**Equation 2: Trophic level (TL)**

$TL_j = 1 + \sum_{i=1}^{S} l_{ij} \frac{TL_i}{n_j}$ (Eq.2)

Where TLj is the trophic level of species j; S is the number of species in the food web; lij is the connection matrix with S rows and S columns, in which for column j and row i, lij = 1 if species j consumes species i and lij = 0 if not; and nj is the number of prey species in the diet of species j. For basal species TL = 1 given that they have no prey. We considered the food web mean TL as the average of all species' TL.

**Equation 3: Modularity (M)**

Modularity is calculated as the difference between realized and expected within-modules interactions, divided by the total number of interactions. We used a stochastic algorithm called “simulated annealing” (Guimera 2005; Saravia et al. 2018), that assumes that the nodes of the same module have more links than one would expect in a random network. The modules are obtained from dividing all the nodes in the network to maximize modularity. In this way, the modularity index is defined as:

$M = \sum_{s} \left( \frac{I_s}{L} - \left( \frac{d_s}{2L} \right)^2 \right)$ (Eq.3)

Where $s$ is the number of modules or compartments, $I_s$ is the number of links between species in the modules, $d_s$ is the sum of degrees for all species in module *s* and *L* is the total number of links.

**Equation 4: Topological roles**

Species topological roles were calculated using the method of functional cartography (Guimera 2005). Roles are determined according to two parameters:

a) the standardized within-module degree (*dz)*, a z-score that reflects how well a species is connected to other species inside the module, relative to other species within its own module:

$$dz_i = \frac{k_{is} - \overline{k_s}}{_{ks}}$$

where $k_{is}$ is the number of links of species $i$ within its own module $s$, $\overline{k_s}$ and $_{ks}$ are the average and standard deviation of $k_{is}$ over all species in $s$, respectively.

b) the participation coefficient (*PC)*, estimates the links distribution of species *i* among modules:

$$PC_i = 1 - \sum_s \frac{k_{is}}{k_i}$$

where $k_i$ is the total number of links of species *i* and $k_{is}$ is the number of links of species *i* to species in module *s*.

To determine the role of each species, the *dz − PC* parameters space was divided into four regions (Kortsch et al. 2015), with two threshold values: *dz* = 2.5 and *PC* = 0.625. Thus, the species were classified as follows:

- *Module hub* (*dz* ≥ 2.5, *PC* < 0.625): The species has a relatively high number of links, but at least 60% within its own module.
- *Module specialist* (*dz* < 2.5, *PC* < 0.625): The species has relatively few links and most within its own module.
- *Module connector* (*dz* < 2.5, *PC* ≥ 0.625): The species has relatively few links and most between modules.
- *Network connector* (*dz* ≥ 2.5, *PC* ≥ 0.625): The species has a high connectivity between and within the modules.

### References

Guimera, Roger. 2005. "Functional Cartography of Complex Metabolic Networks" 433 (February): 895–900. https://doi.org/10.1038/nature03286.1.

Kortsch, Susanne, Raul Primicerio, Maria Fossheim, Andrey V. Dolgov, and Michaela Aschan. 2015. "Climate Change Alters the Structure of Arctic Marine Food Webs Due to Poleward Shifts of Boreal Generalists." *Proceedings of the Royal Society B: Biological Sciences* 282 (1814): 20151546. https://doi.org/10.1098/rspb.2015.1546.

Saravia, Leonardo A, Tomas Ignacio Marina, Marleen De Troch, and Fernando R Momo. 2018. "Ecological Network Assembly: How the Regional Meta Web Influence Local Food Webs." *BioRxiv*, no. June: 340430. https://doi.org/10.1101/340430.

### Supplementary Tables

**Table S1.** Model fit of exponential, log normal, Poisson, power-law, power-law with exponential cutoff and uniform models for degree distributions of Potter Cove and Beagle Channel food webs. AICc and AIC△ are the Akaike corrected for small sample size and delta values for each candidate model. * Indicates best-fit model.

| Food web | Model | AICc | AIC △ |
|---|---|---|---|

| | | | |
|---|---|---|---|
| Potter Cove | Exponential* | 755.51 | 0.00 |
| | Log normal | 768.04 | 12.53 |
| | Poisson | 1296.2 7 | 540.7 6 |
| | Power law | 888.60 | 133.0 9 |
| | Power law /exp | 756.55 | 1.04 |
| | Uniform | 927.16 | 171.6 5 |
| Beagle Channel | Exponential | 1075.0 5 | 0.30 |
| | Log normal | 1089.8 7 | 16.48 |
| | Poisson | 1979.9 6 | 906.5 6 |
| | Power law | 1280.7 3 | 207.3 3 |
| | Power law/exp * | 1073.4 0 | 0.00 |
| | Uniform | 1238.2 9 | 164.8 9 |

**Table S2.** List of trophic species (in increasing trophic level order within module affiliation) included in the Potter Cove food web, their module, topological role, functional group, habitat use, degree (number of trophic interactions) and trophic level (TL). Colors indicate species' topological role: dark purple=network connector (species with high connectivity between and within modules), light purple=module connector (species with few links mostly between modules), light green=module specialist (species with few links within its own module), dark green=module hub (species with high number of links mostly within its own module).

| Trophic species | Module | Topological role | Functional group | Habitat | Degree | TL |
|---|---|---|---|---|---|---|
| Adenocystis utricularis | 1 | Module specialist | basal taxa | benthic | 3 | 1,00 |
| Ascoseira mirabilis | 1 | Module specialist | basal taxa | benthic | 2 | 1,00 |
| Callophyllis atrosanguinea | 1 | Module specialist | basal taxa | benthic | 1 | 1,00 |
| Curdiea racovitzae | 1 | Module specialist | basal taxa | benthic | 2 | 1,00 |
| Desmarestia anceps | 1 | Module specialist | basal taxa | benthic | 6 | 1,00 |
| Desmarestia antarctica | 1 | Module specialist | basal taxa | benthic | 6 | 1,00 |
| Desmarestia menziesii | 1 | Module specialist | basal taxa | benthic | 7 | 1,00 |
| Geminocarpus geminatus | 1 | Module specialist | basal taxa | benthic | 1 | 1,00 |
| Georgiella confluens | 1 | Module specialist | basal taxa | benthic | 3 | 1,00 |

| | | | | | | |
|---|---|---|---|---|---|---|
| Gigartina skottsbergii | 1 | Module specialist | basal taxa | benthic | 8 | 1,00 |
| Iridaea cordata | 1 | Module specialist | basal taxa | benthic | 6 | 1,00 |
| Lambia antarctica | 1 | Module specialist | basal taxa | benthic | 1 | 1,00 |
| Monostroma hariotii | 1 | Module specialist | basal taxa | benthic | 2 | 1,00 |
| Myriogramme manginii | 1 | Module specialist | basal taxa | benthic | 4 | 1,00 |
| Neuroglossum delesseriae | 1 | Module specialist | basal taxa | benthic | 1 | 1,00 |
| Palmaria decipiens | 1 | Module specialist | basal taxa | benthic | 11 | 1,00 |
| Pantoneura plocamioides | 1 | Module specialist | basal taxa | benthic | 1 | 1,00 |
| Phaeurus antarcticus | 1 | Module specialist | basal taxa | benthic | 2 | 1,00 |
| Picconiella plumosa | 1 | Module specialist | basal taxa | benthic | 1 | 1,00 |
| Plocamium cartilagineum | 1 | Module specialist | basal taxa | benthic | 6 | 1,00 |
| Porphyra plocamiestris | 1 | Module specialist | basal taxa | benthic | 1 | 1,00 |
| Trematocarpus antarcticus | 1 | Module specialist | basal taxa | benthic | 1 | 1,00 |
| Ulothrix sp. | 1 | Module specialist | basal taxa | benthic | 1 | 1,00 |
| Urospora penicilliformis | 1 | Module specialist | basal taxa | benthic | 1 | 1,00 |
| Djerboa furcipes | 1 | Module connector | zooplankton | benthic | 8 | 2,00 |
| Eurymera monticulosa | 1 | Module connector | zooplankton | benthic | 8 | 2,00 |
| Gitanopsis squamosa | 1 | Module connector | zooplankton | benthic | 8 | 2,00 |
| Gondogeneia antarctica | 1 | Module specialist | zooplankton | benthic | 20 | 2,00 |
| Laevilacunaria antarctica | 1 | Module specialist | benthos | benthic | 11 | 2,00 |
| Margarella antarctica | 1 | Module connector | benthos | benthic | 5 | 2,00 |
| Nacella concinna | 1 | Module connector | benthos | benthic | 10 | 2,00 |
| Oradarea bidentata | 1 | Module connector | zooplankton | benthic | 7 | 2,00 |
| Paradexamine fissicauda | 1 | Module connector | zooplankton | benthic | 10 | 2,00 |
| Paradexamine sp. | 1 | Module specialist | zooplankton | benthic | 5 | 2,00 |
| Pariphimedia integricauda | 1 | Module specialist | zooplankton | benthic | 8 | 2,00 |
| Probolisca ovata | 1 | Module connector | zooplankton | benthic | 6 | 2,00 |
| Prostebbingia gracilis | 1 | Module specialist | zooplankton | benthic | 17 | 2,00 |
| Prostebbingia sp. | 1 | Module specialist | zooplankton | benthic | 18 | 2,00 |
| Cheirimedon femoratus | 1 | Module connector | zooplankton | benthic | 9 | 2,52 |
| Bovallia gigantea | 1 | Module connector | zooplankton | benthic | 20 | 2,89 |
| Notothenia coriiceps | 1 | Network connector | fish | benthopelagic | 67 | 3,00 |
| Notothenia rossii | 1 | Module connector | fish | benthopelagic | 41 | 3,14 |
| Trematomus newnesi | 1 | Module connector | fish | benthopelagic | 26 | 3,49 |
| Harpagifer antarcticus | 1 | Module connector | fish | benthic | 28 | 3,49 |
| Aged detritus | 2 | Module connector | non-living | benthic | 26 | 1,00 |
| Cumacea | 2 | Module connector | benthos | benthic | 9 | 2,00 |
| Hippomedon kergueleni | 2 | Module connector | zooplankton | benthic | 6 | 2,00 |

| | | | | | | |
|---|---|---|---|---|---|---|
| Oligochaeta | 2 | Module specialist | benthos | benthic | 6 | 2,00 |
| Pseudorchomene plebs | 2 | Module connector | zooplankton | benthic | 4 | 2,00 |
| Spionidae | 2 | Module specialist | benthos | benthic | 5 | 2,00 |
| Gammaridea | 2 | Module connector | zooplankton | benthic | 22 | 2,36 |
| Terebellidae | 2 | Module specialist | benthos | benthic | 6 | 2,45 |
| Nereididae | 2 | Module connector | benthos | benthic | 22 | 2,77 |
| Aglaophamus trissophyllus | 2 | Module connector | benthos | benthic | 9 | 2,97 |
| Nemertea | 2 | Module specialist | benthos | benthic | 14 | 3,15 |
| Polynoidae | 2 | Module connector | benthos | benthic | 18 | 3,19 |
| Priapulida | 2 | Module connector | benthos | benthic | 5 | 3,28 |
| Hydrozoa | 2 | Module connector | benthos | benthic | 19 | 3,29 |
| Barrukia cristata | 2 | Module connector | benthos | benthic | 7 | 3,46 |
| Trematomus bernacchii | 2 | Module connector | fish | benthopelagic | 37 | 3,57 |
| Benthic Diatomea | 3 | Module connector | basal taxa | benthic | 36 | 1,00 |
| Epiphytic Diatomea | 3 | Module specialist | basal taxa | benthic | 7 | 1,00 |
| Fresh detritus | 3 | Module connector | non-living | benthopelagic | 42 | 1,00 |
| Necromass | 3 | Module connector | non-living | benthopelagic | 25 | 1,00 |
| Bryozoa | 3 | Module specialist | benthos | benthic | 11 | 2,00 |
| Chalinidae | 3 | Module specialist | benthos | benthic | 8 | 2,00 |
| Dendrilla antarctica | 3 | Module specialist | benthos | benthic | 3 | 2,00 |
| Laternula elliptica | 3 | Module specialist | benthos | benthic | 9 | 2,00 |
| Porifera | 3 | Module specialist | benthos | benthic | 18 | 2,00 |
| Rosella antartica | 3 | Module specialist | benthos | benthic | 9 | 2,00 |
| Rossella sp. | 3 | Module specialist | benthos | benthic | 8 | 2,00 |
| Stylo_Myca | 3 | Module specialist | benthos | benthic | 11 | 2,00 |
| Tanaidacea | 3 | Module connector | benthos | benthic | 9 | 2,00 |
| Charcotia obesa | 3 | Module connector | zooplankton | benthic | 9 | 2,20 |
| Eatoniella sp. | 3 | Module specialist | benthos | benthic | 6 | 2,33 |
| Polyplacophora | 3 | Module connector | benthos | benthic | 10 | 2,33 |
| Aequiyoldia eightsii | 3 | Module connector | benthos | benthic | 12 | 2,41 |
| Orchomenella sp. | 3 | Module connector | zooplankton | benthic | 11 | 2,43 |
| Neobuccinum eatoni | 3 | Module specialist | benthos | benthic | 9 | 2,50 |
| Doris kerguelenensis | 3 | Module specialist | benthos | benthic | 9 | 2,75 |
| Perknaster fuscus antarticus | 3 | Module specialist | benthos | benthic | 5 | 2,75 |
| Hemiarthrum setulosum | 3 | Module connector | benthos | benthic | 6 | 2,76 |
| Gastropoda | 3 | Module connector | benthos | benthic | 18 | 2,83 |
| Diplasterias brucei | 3 | Module specialist | benthos | benthic | 7 | 2,96 |

| | | | | | | |
|---|---|---|---|---|---|---|
| Perknaster aurorae | 3 | Module specialist | benthos | benthic | 1 | 3,00 |
| Sterechinus neumayeri | 3 | Module connector | benthos | benthic | 21 | 3,00 |
| Odontaster meridionalis | 3 | Module specialist | benthos | benthic | 7 | 3,08 |
| Odontaster validus | 3 | Module specialist | benthos | benthic | 24 | 3,10 |
| Parborlasia corrugatus | 3 | Module specialist | benthos | benthic | 13 | 3,11 |
| Ophionotus victoriae | 3 | Module connector | benthos | benthic | 32 | 3,30 |
| Urticinopsis antarctica | 3 | Module specialist | benthos | benthic | 10 | 3,91 |
| Phytoplankton | 4 | Module connector | basal taxa | pelagic | 23 | 1,00 |
| Ostracoda | 4 | Module connector | zooplankton | benthopelagic | 20 | 2,36 |
| Ascidiacea | 4 | Module connector | benthos | benthic | 9 | 2,45 |
| Copepoda | 4 | Module connector | zooplankton | benthopelagic | 27 | 2,60 |
| Mysida | 4 | Module connector | zooplankton | benthopelagic | 16 | 2,72 |
| Zooplankton | 4 | Module specialist | zooplankton | benthopelagic | 17 | 2,80 |
| Malacobelemnon daytoni | 4 | Module specialist | benthos | benthic | 2 | 2,90 |
| Serolis sp. | 4 | Module connector | benthos | benthic | 7 | 3,03 |
| Polychaeta | 4 | Module connector | benthos | benthic | 27 | 3,06 |
| Salpidae | 4 | Module specialist | zooplankton | pelagic | 13 | 3,28 |
| Gobionotothen gibberifrons | 4 | Module connector | fish | benthic | 15 | 3,32 |
| Glyptonotus antarcticus | 4 | Module connector | benthos | benthic | 15 | 3,37 |
| Euphausia superba | 4 | Module specialist | zooplankton | pelagic | 17 | 3,42 |
| Lindbergichthys nudifrons | 4 | Module connector | fish | benthic | 21 | 3,56 |
| Cephalopoda | 4 | Module specialist | benthos | benthopelagic | 8 | 3,69 |
| Octopoda | 4 | Module connector | benthos | benthopelagic | 8 | 3,75 |
| Hyperiidea | 4 | Module specialist | zooplankton | pelagic | 10 | 3,80 |
| Parachaenichthys charcoti | 4 | Module specialist | fish | benthic | 3 | 4,07 |
| Chaenocephalus aceratus | 4 | Module specialist | fish | benthic | 9 | 4,42 |

**Table S3.** List of trophic species (in increasing trophic level order within module affiliation) included in the Beagle Channel food web, their module, topological role, functional group, habitat use, degree (number of trophic interactions) and trophic level (TL). Colors indicates species' topological role: dark purple=network connector (species with high connectivity between and within modules), light purple=module connector (species with few links mostly between modules), light green=module specialist (species with few links within its own module), dark green=module hub (species with high number of links mostly within its own module).

| Trophic species | Module | Topological role | Functional group | Habitat | Degree | TL |
|---|---|---|---|---|---|---|
| Crustacea | 1 | Module connector | benthos | benthopelagic | 21 | 2,00 |
| Porifera | 1 | Module connector | benthos | benthic | 17 | 2,00 |
| Campylonotus vagans | 1 | Module specialist | benthos | benthic | 4 | 2,00 |
| Ctenophora | 1 | Module specialist | zooplankton | pelagic | 5 | 2,00 |

| | | | | | | |
|---|---|---|---|---|---|---|
| Echiura | 1 | Module specialist | benthos | benthic | 5 | 2,00 |
| Holothuroidea | 1 | Module specialist | benthos | benthic | 6 | 2,00 |
| Careproctus pallidus | 1 | Module specialist | fish | benthic | 1 | 2,10 |
| Copepoda | 1 | Module connector | zooplankton | benthopelagic | 42 | 2,11 |
| Amphipoda | 1 | Module connector | zooplankton | benthopelagic | 40 | 2,20 |
| Euphausiacea | 1 | Module specialist | zooplankton | pelagic | 25 | 2,49 |
| Zygochlamys patagonica | 1 | Module connector | benthos | benthic | 10 | 2,70 |
| Polychaeta | 1 | Module connector | benthos | benthic | 47 | 2,74 |
| Gastropoda | 1 | Module connector | benthos | benthic | 26 | 2,80 |
| Sprattus fuegensis | 1 | Module specialist | fish | pelagic | 17 | 2,88 |
| Galaxias maculatus | 1 | Module specialist | fish | benthopelagic | 6 | 2,89 |
| Ophiuroidea | 1 | Module connector | benthos | benthic | 24 | 2,95 |
| Congiopodus peruvianus | 1 | Module specialist | fish | benthic | 4 | 3,05 |
| Patagonotothen longipes | 1 | Module connector | fish | benthopelagic | 14 | 3,17 |
| Hyperiidea | 1 | Module specialist | zooplankton | pelagic | 11 | 3,18 |
| Stromateus brasiliensis | 1 | Module specialist | fish | benthopelagic | 6 | 3,21 |
| Patagonotothen sp. | 1 | Module specialist | fish | benthopelagic | 9 | 3,24 |
| Patagonotothen sima | 1 | Module specialist | fish | benthopelagic | 7 | 3,29 |
| Odontesthes nigricans | 1 | Module connector | fish | pelagic | 11 | 3,30 |
| Cephalopoda | 1 | Module specialist | benthos | benthopelagic | 23 | 3,35 |
| Merluccius hubbsi | 1 | Module specialist | fish | benthopelagic | 22 | 3,38 |
| Patagonotothen ramsayi | 1 | Module specialist | fish | benthopelagic | 32 | 3,42 |
| Myxine glutinosa | 1 | Module specialist | fish | benthic | 13 | 3,48 |
| Patagonotothen brevicauda brevicauda | 1 | Module specialist | fish | benthic | 8 | 3,51 |
| Priapulida | 1 | Module specialist | benthos | benthic | 8 | 3,53 |
| Schroederichthys bivius | 1 | Module specialist | fish | benthic | 23 | 3,62 |
| Bathyraja albomaculata | 1 | Module specialist | fish | benthic | 19 | 3,63 |
| Macruronus magellanicus | 1 | Module specialist | fish | benthopelagic | 22 | 3,69 |
| Bathyraja scaphiops | 1 | Module specialist | fish | benthic | 6 | 3,70 |
| Bathyraja brachyurops | 1 | Module specialist | fish | benthic | 25 | 3,72 |
| Salilota australis | 1 | Module specialist | fish | benthic | 25 | 3,74 |
| Bathyraja griseocauda | 1 | Module specialist | fish | benthic | 20 | 3,75 |
| Dissostichus eleginoides | 1 | Module specialist | fish | benthopelagic | 28 | 3,77 |
| Genypterus blacodes | 1 | Module specialist | fish | benthic | 33 | 3,78 |
| Cottoperca gobio | 1 | Module specialist | fish | benthic | 31 | 3,83 |
| Raja (Dipturus) chilensis | 1 | Module specialist | fish | benthic | 21 | 3,83 |
| Champsocephalus esox | 1 | Module specialist | fish | benthopelagic | 7 | 3,91 |

| | | | | | | |
|---|---|---|---|---|---|---|
| Diatomea | 2 | Module connector | basal taxa | benthopelagic | 33 | 1,00 |
| Necromass | 2 | Module connector | non-living | benthopelagic | 21 | 1,00 |
| Polysiphonia sp. | 2 | Module connector | basal taxa | benthic | 7 | 1,00 |
| Aged detritus | 2 | Module specialist | non-living | benthic | 19 | 1,00 |
| Desmarestia sp. | 2 | Module specialist | basal taxa | benthic | 3 | 1,00 |
| Porphyra sp. | 2 | Module specialist | basal taxa | benthic | 4 | 1,00 |
| Cumacea | 2 | Module connector | benthos | benthic | 10 | 2,00 |
| Exosphaeroma gigas | 2 | Module connector | benthos | benthic | 11 | 2,00 |
| Tanaidacea | 2 | Module connector | benthos | benthic | 7 | 2,00 |
| Oligochaeta | 2 | Module specialist | benthos | benthic | 5 | 2,00 |
| Laevilitorina caliginosa | 2 | Module connector | benthos | benthic | 15 | 2,09 |
| Terebellidae | 2 | Module connector | benthos | benthic | 8 | 2,11 |
| Gammaridea | 2 | Module specialist | zooplankton | benthic | 28 | 2,13 |
| Cirratulidae | 2 | Module specialist | benthos | benthic | 6 | 2,15 |
| Ostracoda | 2 | Module connector | zooplankton | benthopelagic | 27 | 2,20 |
| Nereididae | 2 | Module specialist | benthos | benthic | 19 | 2,32 |
| Glabraster antarctica | 2 | Module specialist | benthos | benthic | 7 | 2,40 |
| Hydrozoa | 2 | Module connector | benthos | benthic | 26 | 2,65 |
| Glyceridae | 2 | Module specialist | benthos | benthic | 7 | 2,77 |
| Polynoidae | 2 | Module specialist | benthos | benthic | 21 | 2,85 |
| Nemertea | 2 | Module connector | benthos | benthic | 22 | 2,87 |
| Patagonotothen cornucola | 2 | Module connector | fish | benthic | 23 | 3,20 |
| Paranotothenia magellanica | 2 | Module connector | fish | benthopelagic | 14 | 3,28 |
| Harpagifer bispinis | 2 | Module connector | fish | benthic | 15 | 3,30 |
| Fresh detritus | 3 | Module connector | non-living | benthopelagic | 45 | 1,00 |
| Macroalgae | 3 | Module connector | basal taxa | benthic | 21 | 1,00 |
| Phytoplankton | 3 | Module connector | basal taxa | pelagic | 42 | 1,00 |
| Blidingia minima | 3 | Module specialist | basal taxa | benthic | 1 | 1,00 |
| Cladophora sp. | 3 | Module specialist | basal taxa | benthic | 6 | 1,00 |
| Codium sp. | 3 | Module specialist | basal taxa | benthic | 3 | 1,00 |
| Derbesia sp. | 3 | Module specialist | basal taxa | benthic | 1 | 1,00 |
| Ulothrix sp. | 3 | Module specialist | basal taxa | benthic | 1 | 1,00 |
| Bryozoa | 3 | Module connector | benthos | benthic | 14 | 2,00 |
| Bivalvia | 3 | Module specialist | benthos | benthic | 19 | 2,00 |
| Calliostoma nudum | 3 | Module specialist | benthos | benthic | 7 | 2,00 |
| Chthamalus sp. | 3 | Module specialist | benthos | benthic | 4 | 2,00 |
| Eurhomalea exalbida | 3 | Module specialist | benthos | benthic | 2 | 2,00 |
| Fissurella oriens | 3 | Module specialist | benthos | benthic | 4 | 2,00 |

| | | | | | | |
|---|---|---|---|---|---|---|
| Gaimardia trapesina | 3 | Module specialist | benthos | benthic | 8 | 2,00 |
| Hiatella arctica | 3 | Module specialist | benthos | benthic | 7 | 2,00 |
| Margarella violacea | 3 | Module specialist | benthos | benthic | 13 | 2,00 |
| Membranipora isabelleana | 3 | Module specialist | benthos | benthic | 10 | 2,00 |
| Nacella mytilina | 3 | Module specialist | benthos | benthic | 5 | 2,00 |
| Notobalanus flosculus | 3 | Module specialist | benthos | benthic | 4 | 2,00 |
| Pareuthria fuscata | 3 | Module specialist | benthos | benthic | 7 | 2,00 |
| Plaxiphora sp. | 3 | Module specialist | benthos | benthic | 5 | 2,00 |
| Serpulidae | 3 | Module specialist | benthos | benthic | 7 | 2,00 |
| Spirorbinae | 3 | Module specialist | benthos | benthic | 8 | 2,00 |
| Zooplankton | 3 | Module connector | zooplankton | benthopelagic | 42 | 2,02 |
| Fissurella picta | 3 | Module specialist | benthos | benthic | 13 | 2,13 |
| Crepipatella dilatata | 3 | Module specialist | benthos | benthic | 9 | 2,15 |
| Mytilus edulis chilensis | 3 | Module specialist | benthos | benthic | 21 | 2,16 |
| Ascidiacea | 3 | Module specialist | benthos | benthic | 8 | 2,16 |
| Aulacomya atra | 3 | Module specialist | benthos | benthic | 15 | 2,20 |
| Cirripedia | 3 | Module specialist | benthos | benthic | 6 | 2,25 |
| Perumytilus purpuratus | 3 | Module specialist | benthos | benthic | 8 | 2,25 |
| Notochthamalus scabrosus | 3 | Module specialist | benthos | benthic | 7 | 2,34 |
| Halicarcinus planatus | 3 | Module connector | benthos | benthic | 26 | 2,42 |
| Pseudechinus magellanicus | 3 | Module connector | benthos | benthic | 45 | 2,58 |
| Tonicia sp. | 3 | Module specialist | benthos | benthic | 6 | 2,67 |
| Pagurus comptus | 3 | Module connector | benthos | benthic | 27 | 2,85 |
| Arbacia dufresnii | 3 | Module specialist | benthos | benthic | 23 | 2,85 |
| Acanthocyclus albatrossis | 3 | Module connector | benthos | benthic | 11 | 2,89 |
| Paralomis granulosa | 3 | Module specialist | benthos | benthic | 26 | 3,16 |
| Peltarion spinosulum | 3 | Module specialist | benthos | benthic | 25 | 3,16 |
| Trophon geversianus | 3 | Module specialist | benthos | benthic | 16 | 3,19 |
| Lithodes santolla | 3 | Module specialist | benthos | benthic | 30 | 3,22 |
| Eurypodius latreillii | 3 | Module specialist | benthos | benthic | 25 | 3,23 |
| Anasterias antarctica | 3 | Module specialist | benthos | benthic | 39 | 3,25 |
| Asterina fimbriata | 3 | Module specialist | benthos | benthic | 14 | 3,26 |
| Cosmasterias lurida | 3 | Module specialist | benthos | benthic | 25 | 3,31 |
| Ulva sp. | 4 | Module connector | basal taxa | benthic | 17 | 1,00 |
| Adenocystis utricularis | 4 | Module specialist | basal taxa | benthic | 9 | 1,00 |
| Ballia sp. | 4 | Module specialist | basal taxa | benthic | 3 | 1,00 |
| Bostrychia sp. | 4 | Module specialist | basal taxa | benthic | 4 | 1,00 |
| Callophyllis pinnata | 4 | Module specialist | basal taxa | benthic | 7 | 1,00 |

| | | | | | | |
|---|---|---|---|---|---|---|
| Ceramium diaphanum | 4 | Module specialist | basal taxa | benthic | 8 | 1,00 |
| Ceramium sp. | 4 | Module specialist | basal taxa | benthic | 5 | 1,00 |
| Delesseriaceae | 4 | Module specialist | basal taxa | benthic | 3 | 1,00 |
| Ectocarpus sp. | 4 | Module specialist | basal taxa | benthic | 6 | 1,00 |
| Griffithsia sp. | 4 | Module specialist | basal taxa | benthic | 1 | 1,00 |
| Halopteris sp. | 4 | Module specialist | basal taxa | benthic | 4 | 1,00 |
| Hincksia sp. | 4 | Module specialist | basal taxa | benthic | 2 | 1,00 |
| Hymenena sp. | 4 | Module specialist | basal taxa | benthic | 1 | 1,00 |
| Macrocystis pyrifera | 4 | Module specialist | basal taxa | benthic | 21 | 1,00 |
| Monostroma sp. | 4 | Module specialist | basal taxa | benthic | 1 | 1,00 |
| Myriogramme sp. | 4 | Module specialist | basal taxa | benthic | 1 | 1,00 |
| Nothogenia fastigiata | 4 | Module specialist | basal taxa | benthic | 8 | 1,00 |
| Rhizoclonium sp. | 4 | Module specialist | basal taxa | benthic | 6 | 1,00 |
| Scytothamus fasciculatus | 4 | Module specialist | basal taxa | benthic | 7 | 1,00 |
| Sphacelaria sp. | 4 | Module specialist | basal taxa | benthic | 2 | 1,00 |
| Trailliella sp. | 4 | Module specialist | basal taxa | benthic | 1 | 1,00 |
| Ulva rigida | 4 | Module specialist | basal taxa | benthic | 7 | 1,00 |
| Isopoda | 4 | Module connector | benthos | benthic | 41 | 2,00 |
| Siphonaria lessonii | 4 | Module specialist | benthos | benthic | 18 | 2,00 |
| Loxechinus albus | 4 | Module specialist | benthos | benthic | 5 | 2,18 |
| Polyplacophora | 4 | Module specialist | benthos | benthic | 23 | 2,19 |
| Nacella deaurata | 4 | Module specialist | benthos | benthic | 30 | 2,27 |
| Munida gregaria | 4 | Network connector | zooplankton | benthopelagic | 71 | 2,37 |
| Nacella magellanica | 4 | Module connector | benthos | benthic | 22 | 2,43 |
| Eleginops maclovinus | 4 | Module connector | fish | benthic | 42 | 3,02 |
| Xymenopsis muriciformis | 4 | Module specialist | benthos | benthic | 9 | 3,21 |
| Patagonotothen tessellata | 4 | Module connector | fish | benthopelagic | 35 | 3,23 |
| Austrolycus depressiceps | 4 | Module connector | fish | benthic | 17 | 3,50 |

**Table S4.** Kolmogorov-Smirnov test D and p values for Potter Cove and Beagle Channel metrics comparison of simulated values. * for p-values significantly different (<0.05).

| Network metric | D | p |
|---|---|---|
| Mean trophic level | 0.99 | <0.01* |
| Omnivory | 1 | <0.01* |
| Modularity | 0.97 | <0.01* |
| QSS | 0.93 | <0.01* |
